# High-throughput single-particle tracking reveals nested membrane nanodomain organization that dictates Ras diffusion and trafficking

**DOI:** 10.1101/552075

**Authors:** Yerim Lee, Carey Phelps, Tao Huang, Barmak Mostofian, Lei Wu, Ying Zhang, Young Hwan Chang, Philip J. S. Stork, Joe W. Gray, Daniel M. Zuckerman, Xiaolin Nan

## Abstract

Membrane nanodomains have been implicated in Ras signaling, but what these domains are and how they interact with Ras remain obscure. Using high throughput single particle tracking with photoactivated localization microscopy and detailed trajectory analysis, here we show that distinct membrane domains dictate KRas diffusion and trafficking in U2OS cells. KRas exhibits an immobile state in domains ∼70 nm in size, each embedded in a larger domain (∼200 nm) that confers intermediate mobility, while the rest of the membrane supports fast diffusion. Moreover, KRas is continuously removed from the membrane via the immobile state and replenished to the fast state, likely coupled to internalization and recycling. Importantly, both the diffusion and trafficking properties of KRas remain invariant over a broad range of protein expression levels. Our results reveal how membrane organization dictates KRas diffusion and trafficking and offer insight into how Ras signaling may be regulated through spatial mechanisms.

## Introduction

The plasma membrane has a complex and dynamic landscape that helps shape how diverse membrane-localized signaling molecules behave^1–6^. Among others, the Ras small GTPases are prototypical examples of signaling molecules whose biological activities are directly regulated by the membrane^7,8^. While biochemical aspects of how Ras interacts with downstream effectors such as Raf have been well studied^9,10^, the mechanisms through which the biological membrane defines the signaling activity and specificity of Ras are still poorly understood. Recent studies by us and others suggest that Ras signaling may involve the formation of multimers (dimers and/or clusters) in a membrane-dependent manner^11–17^, and that partitioning of Ras into nanoscopic membrane domains and interactions with scaffold proteins or structures likely constitute critical steps to Ras multimer formation and signaling^18–22^. While previous high-resolution imaging experiments using immuno-EM^15,17^ or quantitative superresolution microscopy^12,23^ were instrumental to revealing the existence of Ras multimers, the resulting images were mostly static and provided limited information about the spatiotemporal dynamics of Ras – membrane domain interactions.

Live-cell single-particle tracking (SPT)^24–26^ complements static imaging by providing information about molecular motions, and it has been used to study Ras dynamics on the membrane^27–29^. The underlying rationale is that interactions of Ras with different membrane domains and signaling partners would manifest as varying diffusion behavior. Indeed, using SPT, Murakoshi *et al.* observed transient events of Ras immobilization on the membrane, which became more frequent upon epidermal growth factor stimulation, potentially reflecting the formation of signaling complexes or interactions with raft domains^28^. Lommerse and colleagues also used SPT to probe Ras diffusion and similarly observed transient and context-dependent confinement of Ras in membrane regions not more than 200 nm in diameter^27^.

These prior studies offered important initial insight into the potential connections between Ras diffusion, function, and membrane organization, but the technical constraints of traditional SPT resulted in low throughput which limited the depth of these studies. Typically, only a few tens of trajectories could be obtained from each experiment, which precluded detailed and quantitative characterization of the heterogeneous and stochastic nature of molecular diffusion. In consequence, while the studies consistently reported two diffusion states – a ‘free’ diffusion state and another ‘immobile’ state, it remains to be seen whether a two-state model accurately recapitulates Ras membrane dynamics^27–29^. Thus, the nature of the membrane domains occupied by each of these states and how Ras molecules transition between the states in connection with multimer formation and signaling remain unclear.

Recent years have seen significant advances in both experimental^30–35^ and data analysis strategies^36–44^ of SPT, some of which have dramatically improved the information throughput. Among others, spt-PALM combines SPT with photoactivated localization microscopy (PALM) to enable single molecule tracking under dense labeling conditions through stochastic photoswitching^30^. With spt-PALM, it is routine to acquire tens of thousands of diffusion trajectories from a single cell. A growing list of software tools has also been developed to facilitate spt-PALM data analysis^36,37,39,43,45^. In particular, variational Bayes SPT (vbSPT) package allows construction of a detailed diffusion model from spt-PALM data with parameters such as the number of states, the diffusion coefficient and the occupancy of each state, as well as the state transition rates even when the individual trajectories are short^36^. Additionally, a wide range of analytical methods has been introduced to gain further insight into the states and the state transitions from SPT trajectories^40,43,46^. These advances help overcome the limitations of conventional SPT and make it possible to analyze Ras membrane dynamics in much greater depth.

Here, we report our efforts on combining spt-PALM with detailed trajectory analysis to reveal previously unknown aspects of Ras diffusion on the cell membrane. With carefully controlled expression levels and photoactivation rate, spt-PALM trajectories of PAmCherry1-tagged KRas G12D consistently reported three diffusion states, including a fast diffusion state, an immobile state, and a previously unidentified diffusion state with intermediate mobility. Leveraging the large number of trajectories, we were able to spatially map the diffusion states to distinctive membrane domains, estimate the size and lifetime of each domain, and define the spatial relationship between the domains. Moreover, in analyzing how KRas transitions from one diffusion state to another, we discovered that KRas diffusion follows a non-equilibrium steady state (NESS) model with net mass flow from the fast state to the immobile state, likely coupled to the endocytic trafficking and membrane recycling of KRas. Based on these results, we propose a new model to describe the membrane dynamics of KRas, where nested membrane nanodomains dictate the diffusion, trafficking, and potentially multimer formation and signaling.

## Results

### KRas diffuses on the membrane in three distinct states

To investigate the lateral diffusion properties of KRas under controlled expression levels, we established isogenic U2OS cells stably expressing PAmCherry1-KRas G12D under doxycycline (Dox) regulation^12^. The expression level of PAmCherry1-KRas G12D was comparable to that of the endogenous KRas at 2 ng/mL Dox as judged by western blotting, with higher expression levels achieved at 5-10 ng/mL Dox (Fig. 1A). Hence, initially data were collected from cells induced at 2 ng/mL Dox. The photoactivatable fluorescent protein PAmCherry1 has been widely used for quantitative PALM and spt-PALM^47^. Owing to the good single-molecule brightness of activated PAmCherry1, we were able to track individual PAmCherry1-KRas G12D molecules at frame rates up to ∼83 Hz (i.e., ∼12 ms/frame) with a low excitation dose (∼400 W/cm^2^ at 561 nm). The low spontaneous photoactivation rate of PAmCherry1 also permits clean single-molecule imaging even at high expression levels, yielding as many as hundreds of thousands of trajectories per cell via spt-PALM (Fig. 1B and Suppl. Video 1).

**Figure 1.**
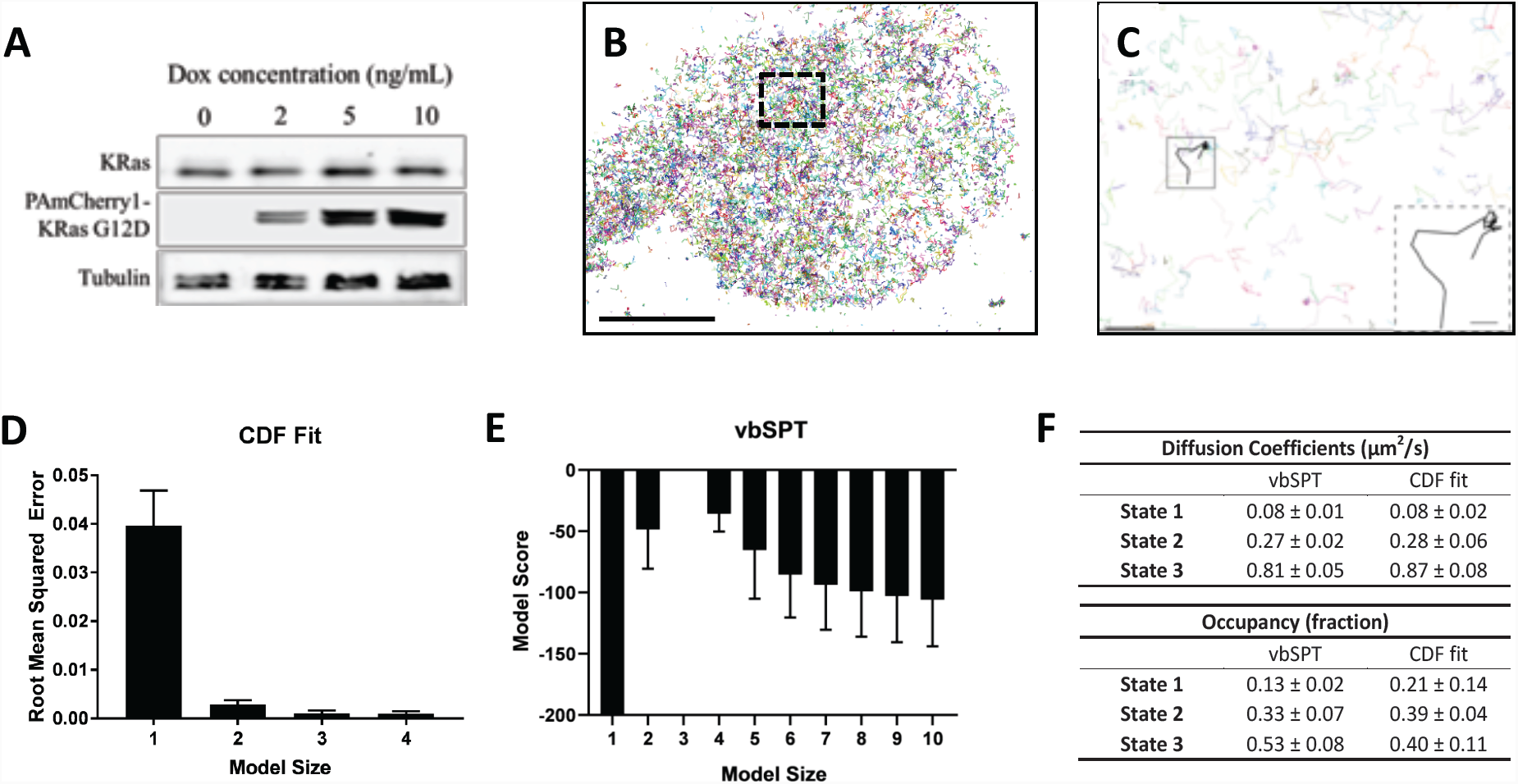
Defining the membrane diffusion state model of KRas using spt-PALM. A) Western blot showing the increasing expression level of PAmCherry1-KRas G12D with increasing doxycycline (Dox) concentration; B) Example trajectory map of membrane KRas acquired at 12 ms frame rate using TIRF illumination. Each line represents an individual Ras molecule coordinate over time acquired for the duration of the movie (20 minutes). Only a subset of all trajectories is plotted. Scale bar: 10 μm; C) Expanded view of the boxed region in B). Only a subset of all of the trajectories in the boxed region is shown to allow unhindered view of individual Ras trajectories. Inset shows a KRas trajectory displaying multiple diffusion states. Scale bars: main figure: 1 mm; inset: 200 nm; D) Determining the optimal model size for KRas membrane diffusion using CDF fitting, with smaller root mean squared error indicating a better model. Error bars are 95% confidence intervals (CIs); E) Determining the optimal model size for KRas membrane diffusion using vbSPT, with smaller absolute model score (i.e., score of zero being the best global model) indicating a better model. Error bars represent 95% CIs; F) Comparing the model parameters obtained from CDF fit and vbSPT, both using a three-state model for KRas membrane diffusion. State transition probabilities were not inferred from CDF fit and therefore not included in the comparison. Error bars are 95% CIs.

A close inspection of the individual trajectories clearly shows larger diffusive steps intermittent with moments of transient entrapment, indicating the presence of multiple diffusion states and frequent state transitions (Fig. 1C and inset). Similar observations were reported for both HRas and KRas in previous low throughput SPT experiments, where two diffusion states – a ‘fast’ state and an ‘immobile’ diffusion state – were detected^27,28^.

With a high frame rate and large number of trajectories, we first probed the number of diffusion states. To this end, we used two methods to analyze the Ras diffusion trajectories. The first approach fits a cumulative distribution function (CDF) for Brownian motion to the squared displacements of Ras trajectories to extract diffusion coefficients and their respective occupancies^48^. The second method, vbSPT, treats particle diffusion and the associated state transitions with a Hidden Markov Model and performs model selection through variational inference^36^.

Both CDF fitting and vbSPT yielded similar three-state models for KRas G12D diffusion on the membrane of live U2OS cells. Specifically, CDF fitting to a three-state model had significantly lower residual error compared to a single- or a two-state model and further increasing the model size did not decrease the error (Fig. 1D), indicating that a three-state model is sufficient to describe the data. For vbSPT, a score equal to zero indicates the best model, a condition that was met with a three-state model but not with larger or smaller size models (Fig. 1E). The diffusion coefficient and the occupancy for each of the diffusion states were in good agreement between the two analysis methods and within each method when applied to different cells under the same conditions, as evidenced by the small errors (Fig. 1F). Datasets with high particle densities can return models with variable sizes, sometimes also with aberrant model parameters (Suppl. Figs. 1B-D and Suppl. Fig. 3); even so, the histogram of all vbSPT-derived diffusion coefficients still showed three distinct clusters (Suppl. Fig. 3C) corresponding to the three states listed in Fig. 1F and Suppl. Fig. 4A. Thus, we concluded that the membrane diffusion of KRas G12D under our experimental conditions is best described by a three-state model, demonstrating the existence of an intermediate state not detected in previous studies. Because vbSPT supplies the transition probabilities and state identities for every time step, vbSPT was used for most subsequent analyses in the remainder of this work.

The diffusion coefficient of the slowest state in Fig. 1F is comparable to that expected from single-molecule localization error (∼30 nm, Suppl. Fig. 5), which implied that the actual diffusion of KRas in this state may be much slower than it appeared. To test this hypothesis, we acquired spt-PALM data at a slower frame rate (35 ms/frame) to improve the localization accuracy of slow moving molecules since more photons could now be collected for each PAmCherry1 molecule in a single frame (Suppl. Fig. Indeed, these datasets reported a significantly smaller diffusion coefficient (0.02 µm^2^/s) for the slowest state than that obtained earlier (0.08 µm^2^/s) using data taken at 12 ms/frame. This result suggests that the slowest diffusion state of KRas is essentially an immobile state, consistent with previous reports^27,28^.

### KRas diffusion states correspond to distinct membrane domains

The diffusion model presented in Fig. 2A summarizes the results from the spt-PALM trajectory analyses above. Each circle represents one of the diffusion states with arrows indicating the transition probabilities between pairs of states. A notable feature of this model is that there appears to be a defined state transition path: KRas molecules always transition between the fast (F) and the immobile (I) states by going through the intermediate (N) state, and direct transitions between the fast and the immobile states almost never occur. In order to confirm this transition path, we compared the distribution of step sizes relative to the immobile state steps, since different step sizes would reflect different diffusion coefficients. In support of this hypothesis, the histogram of step sizes immediately adjacent to the immobile steps corresponded to the intermediate diffusion state (Fig. 2B, blue) while the distribution of the remaining steps had a broader peak implying a mixture of both fast and intermediate diffusion steps (Fig. 2B, where the black color indicates a mixture of states). As expected, the step sizes assigned to the immobile states (Fig. 2B, red) are even smaller compared to that of the other two states. The clear separation of these three step size distributions confirms the above-mentioned transition path through the intermediate state. The distinctions in step sizes among the three states were even more obvious on data taken at 35 ms/frame, which had better single-molecule localization precision (Suppl. Fig. 6A). Thus, the intermediate state is not merely a state with intermediate mobility but effectively an obligatory intermediate between the immobile and the fast states of KRas.

**Figure 2.**
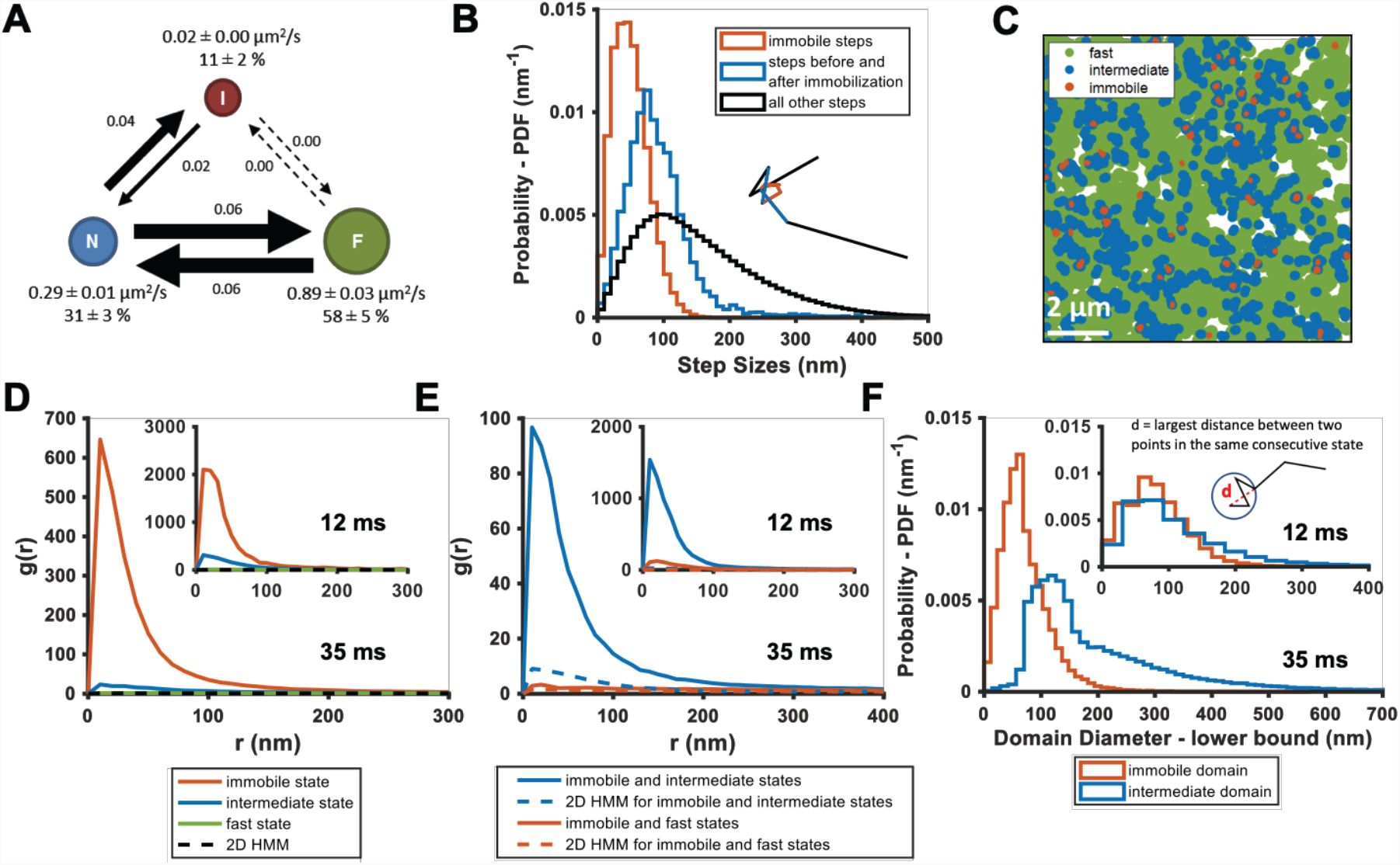
KRas diffusion states are associated with distinct membrane domains. A) The three-state model for KRas diffusion with F, N, and I, representing the fast, the intermediate, and the immobile states, respectively. Model parameters were inferred using vbSPT on spt-PALM datasets with at least 30,000 trajectories obtained on cells induced with 2 ng/mL Dox. The arrows indicate state transitions (i.e. the probability of switching to a different state in the next frame) and the area of the circle and the thickness of the arrows are both roughly scaled to reflect their relative values. All parameters were derived from data acquired at 12 ms frame interval except for the diffusion coefficient of the immobile state, which was inferred from data taken at 35 ms frame interval. Error bars are 95% CIs; B) Step size histograms for immobilization events (red), one step before or after the immobilization event (blue), and all other steps (black). A diffusion step was part of an immobilization event if immobile state was assigned to that trajectory segment by vbSPT (see *Methods*); C) Map of the membrane locations where KRas molecules exhibit specific diffusion states (referred to as state coordinates) within a one-minute duration (taken from a spt-PALM dataset of ∼20 min total duration). Red, blue, and green dots represent locations of the immobile, the intermediate, and the fast states, respectively, with each rendered circles scaled proportionally to the mean diffusion coefficient for the state; D) Pair correlation analysis on the averaged state coordinates across multiple, one-minute segments of longer spt-PALM datasets. The same color coding as in B) was used to distinguish the three states. For this analysis, molecules in the same diffusion state in successive frames only contributed a single, averaged state coordinate. The average state coordinates of all molecules captured within a one-minute segment were used for correlation analysis, and the results from multiple one-minute segments were averaged to yield the plot. The negative control was generated through a 2D Markovian simulation, and the resulting trajectories were analyzed the same as the experiment (see *Methods*); E) Cross correlation analysis between pairs of diffusion states. The state coordinates were processed the same way as in D) prior to the correlation analysis, except that the correlation was performed between two different diffusion states. The negative control was generated through a 2D Markovian simulation, and the resulting trajectories were analyzed the same as the experiment (see *Methods*); F) Estimating the lower bound size for the immobile and the intermediate domains. The estimation was based on the maximum distance traveled by the molecule while in the same diffusion state. *D-F) The main panel shows results inferred from data taken at 35 ms frame intervals for improved localization precision. The inset shows the data taken at 12 ms/frame.

The observed state transition path may arise from at least two potential scenarios. In the first scenario, fast diffusing KRas may spontaneously transition into the intermediate then the immobile state through conformational changes. In this case, the intermediate and the immobile states of KRas would be temporally and spatially correlated, but the states would appear randomly on the membrane. Alternatively, the immobile states could be caused by KRas transiently binding to structures (‘immobilization sites’) residing in specialized membrane regions that confer intermediate mobility to KRas (thus termed ‘intermediate domains’). Consequently, these intermediate domains would act as transition zones between regions of the membrane where KRas exhibits fast diffusion and the KRas immobilization sites, generating the observed state transition path. If the latter were true, there should be a temporal and spatial correlation not only between the intermediate and the immobile states, but also within each of the two states of KRas – that is, the locations for each of the two states should self-cluster, provided that the intermediate domains and immobilization sites have lifetimes longer than our temporal resolution. In other words, we should be able to observe multiple visits to the same intermediate or immobilization domains by different KRas molecules within the lifetime of these domains.

To distinguish between the two scenarios, we performed auto- and cross-correlation analysis on the locations of KRas exhibiting a certain diffusion state (referred to hereafter as state coordinates). We first visually examined the spatial distributions of the states by slicing each raw image stacks into one-minute time segments and plotting the state coordinates on the same map, with each color representing one of the states (Fig. 2C, Suppl. Fig. 7 and Suppl. Video 2). Each diffusion trajectory typically contributes only a few points to the plots as limited by its short duration, and the points from multiple trajectories accumulate over time (up to 1 min in this case) to ‘paint’ a map of the membrane regions associated with each of the diffusion states. As shown in Fig. 2C, the intermediate states and the immobile states not only co-clustered, but also each appeared to self-cluster. Specifically, regions corresponding to the intermediate states (blue) often connect to give rise to nanoscopic domains a few hundred nm in size and the vast majority of the immobilization sites (red) are surrounded by the intermediate domains. By contrast, regions corresponding to the fast state occupy the majority of the membrane area. While both the intermediate and the immobile domains appeared to be dynamic, a time-lapse domain map (Suppl. Video 2) showed that at least some of these domains could last a few minutes (addressed in greater detail in Figure 3). Thus, spatial mapping of the KRas state coordinates provided visual evidence for the physical presence of nested, nanoscopic domains conferring the distinct KRas diffusion states.

**Figure 3.**
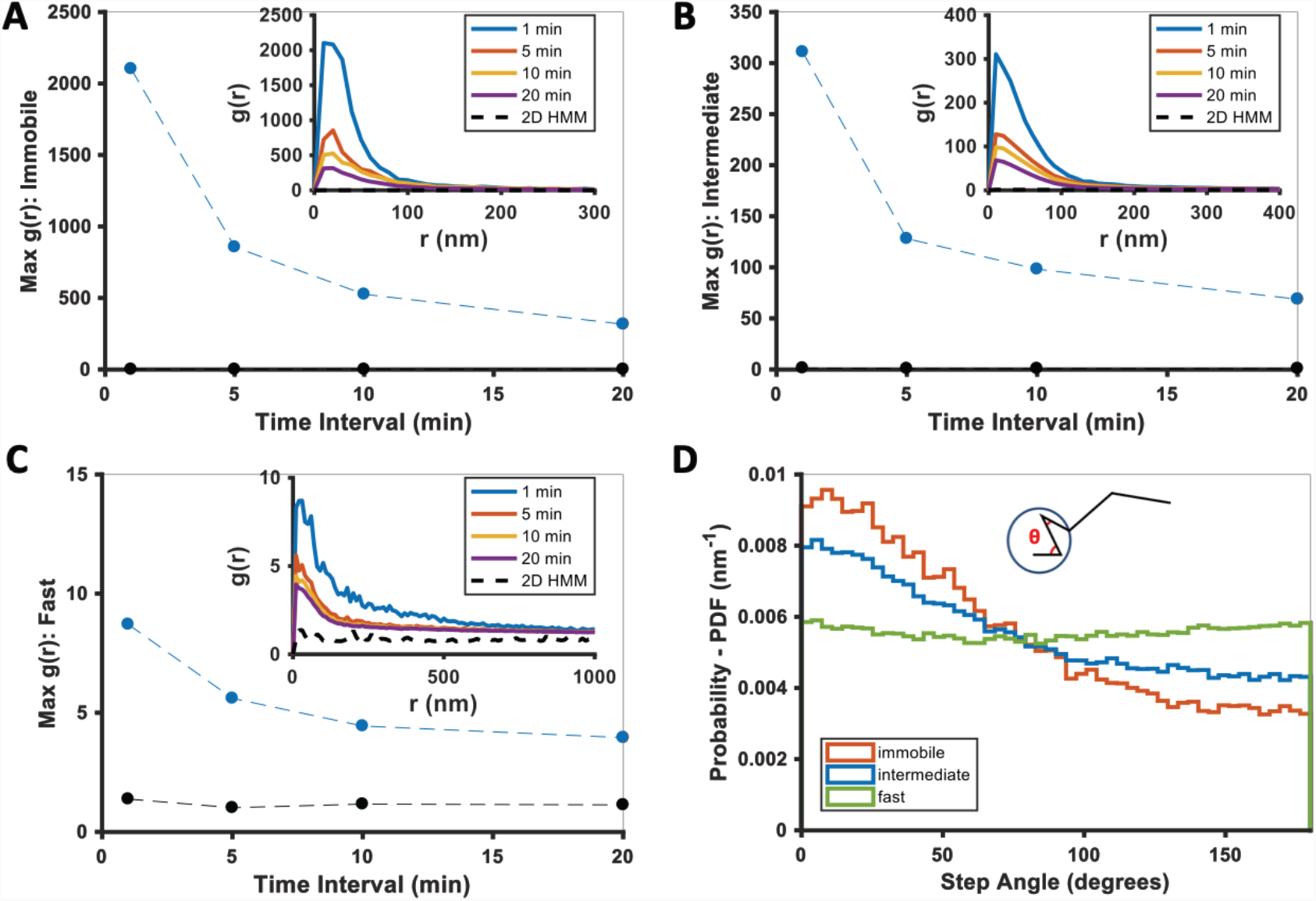
Temporal properties of the KRas-associated immobile and intermediate domains. A-C) Pair correlation analysis of the state coordinates at different time intervals (1, 5, 10, and 20 min). The amplitude (maximal *g(r)* value) of the pair correlation function at each time interval was plotted in the main panel with the raw pair correlation plots shown in the inset. A-C show pair correlation functions of averaged coordinates for the immobile, the intermediate, and the fast states, respectively (see *Methods*). The negative control in each case was generated through a 2D Markovian simulation, and the resulting trajectories were analyzed the same as the experiment (see *Methods*); D) Deflection angle analysis on KRas diffusion trajectories separated by diffusion states (red: immobile; blue: intermediate; green: fast). The deflection angle was calculated as the angle between two successive segments of the trajectory while the molecule was in the same diffusion state. *Results shown for data acquired at 12 ms/frame.

We next used pair correlation function (*g(r)*) to quantitate the spatial relationship between the KRas states. The function *g(r)* measures the ratio of the number of particles located a distance (*r*) from a given particle to that expected from a complete spatial randomness (see *Methods*). Here, the *g(r)* could be calculated for particles in the same diffusion state (auto-correlation) or between two different diffusion states (cross-correlation); in either case, amplitudes of *g(r)* significantly greater than that expected for a random distribution indicate spatial clustering. When multiple KRas molecules visit the same domain, each at a different time point but exhibiting the same diffusive state, *g(r)* would detect spatial auto-correlation for the given state. To avoid false clustering due to the same molecule staying in the same state across multiple frames, we used the averaged state coordinate for each continuous trajectory segment that stayed in the same state for more than two consecutive time points (see *Methods*). Since datasets acquired at 35 ms/frame had significantly improved localization precision (Suppl. Fig. 5), the main panel for all spatial analysis shows results from the 35 ms/frame dataset while the results from data acquired at 12 ms/frame are shown in the inset.

Consistent with the visual observation earlier (Fig. 2C), coordinates of the immobile and the intermediate states each showed significant clustering in the *g(r)* plots averaged across each 1-minute raw image stacks, whereas *g(r)* of the fast state was barely above random across the full range of *r* analyzed (Fig. 2D). All *g(r)* negative controls were generated with a 2D Markovian simulation of diffusing particles with no associated domains (see *Methods*), and the simulated trajectories were processed through the same analysis pipeline as the experimental data. As expected, the averaged state coordinates of the simulated negative control had values close to one and showed no peak in the *g(r)* plots. Furthermore, *g(r)* based on spatial cross-correlation analysis clearly indicated co-clustering between the immobile and the intermediate state positions but not with the fast diffusion state (Fig. 2E). The spatial analysis for trajectories collected from the 35 ms acquisition time is discussed in greater detail in Supplementary Figure 6.

### Transient, nanoscopic domains mediate the intermediate and the immobile states of KRas

We estimated the lower-bound size of the membrane domains associated with the immobile and the intermediate states of KRas by calculating the maximum distance a particle traveled within a domain (i.e., longest distance between two points within consecutive steps taken while in the same state). For this analysis, we focus on the values acquired from 35 ms/frame dataset not only for the improved localization precision (Suppl. Fig. 5) but also for the larger membrane area covered by each molecule in a single step(35 ms vs 12 ms). Shown in Fig. 2F main panel are the histograms of the estimated domain sizes determined from the lower frame rate (35 ms/frame), based on which we determined that the mean diameters of the intermediate and the immobile membrane domains were at least ∼200 nm and ∼70 nm, respectively. This is consistent with the notion that most immobile domains are likely surrounded by intermediate domains. We further note that the distribution of the minimum intermediate domain size appeared to have at least two peaks at ∼120 nm and ∼230 nm, implying that there are potentially multiple types of intermediate domains (Fig. 2F).

In order to further investigate the temporal behavior of the immobile and the intermediate domains, we extended *g(r)* calculations as in Figure 2 from one minute to longer time intervals. The expectation was that, as the time interval for *g(r)* increases beyond the lifetime of a domain, the chance of observing KRas molecules visiting the same nanodomain (i.e., exhibiting the same diffusion state in close proximity) should decrease, resulting in lower *g(r)*. Indeed, as shown in Figure 3A-C, for dataset acquired at 12 ms frame interval, the peak amplitudes of *g(r)* for both the immobile (Fig. 3A) and the intermediate (Fig. 3B) states decreased significantly after ∼5 min with further decay at increasing time intervals, indicative of finite lifetimes for both nanodomains, likely on the order of minutes on average (see also Suppl. Fig. 6D-F for results with data taken at 35 ms/frame). We note that, for the limited temporal resolution, this analysis will only capture relatively stable domains with lifetimes longer than 1 min, and we cannot rule out the presence of other, more transient intermediate or immobile domains.

To gain insight into how KRas interacts with the different membrane domains, we also analyzed the frame-to-frame deflection angle for KRas molecules within each domain. The deflection angle measures the relationship between the current and the preceding step: a complete random walk would yield a flat distribution of deflection angles, whereas a preference for acute angles indicates more ‘returning’ steps. As shown in Fig. 3D, KRas molecules trapped in either the immobile or the intermediate domains (the red and the blue lines) were more likely to exhibit acute deflection angles, potentially due to backward movements at the domain boundaries. In comparison, KRas molecules in the fast state exhibit (Fig. 3D, the green line) equal probabilities of moving in all directions, consistent with Brownian motion.

### KRas is constitutively depleted from the immobile state and replenished to the fast state

The small variance in the estimated model parameters from different cells, either from the same or from different samples (Fig. 2A), led us to hypothesize that KRas membrane diffusion is in a steady state. To verify this, we divided each spt-PALM dataset with a minimum of 40,000 trajectories into four quarters (where each quarter is ∼10,000 trajectories and typically ∼5 min in duration) and computed the diffusion model for each quarter using vbSPT. As Fig. 4A shows, the model parameters for all four quarters were essentially identical, which is the case for all qualifying datasets, confirming that KRas diffusion on the membrane of U2OS cells is indeed in a steady state, at least on the investigated time scales (up to ∼20 minutes).

While the above analysis indicates that the three-state KRas diffusion system should be stable over time, we found that the model as presented in Figure 2A is not self-sustaining. When using experimentally derived model parameters to simulate how the three-state system evolves over time (see *Methods*), we observed that the system quickly deviated from its initial configuration and instead stabilized at an entirely different set of state occupancies (Fig. 4B). In the new, ‘equilibrated’ system configuration, KRas spent as much as ∼50% of its time in the immobile state, which is significantly more than the observed steady state occupancy of ∼11%. The fast state was the opposite; the population residing in the fast state would be significantly reduced from 58% to ∼25%. By contrast, the intermediate state changed only slightly (∼31% vs ∼24% for the experimental and the theoretical observations, respectively). We confirmed that the simulated equilibrium probabilities were consistent with the principle of detailed balance^49^ (Fig. 4C), and further verified that the non-equilibrium occupancies of the observed KRas diffusion model were not an artifact of vbSPT since vbSPT correctly retrieved the steady state model parameters when applied to trajectories generated from steady state simulations with different input parameters (Suppl. Fig. 8). Therefore, we concluded that the model in Fig. 2A represents a non-equilibrium steady state (NESS).

**Figure 4.**
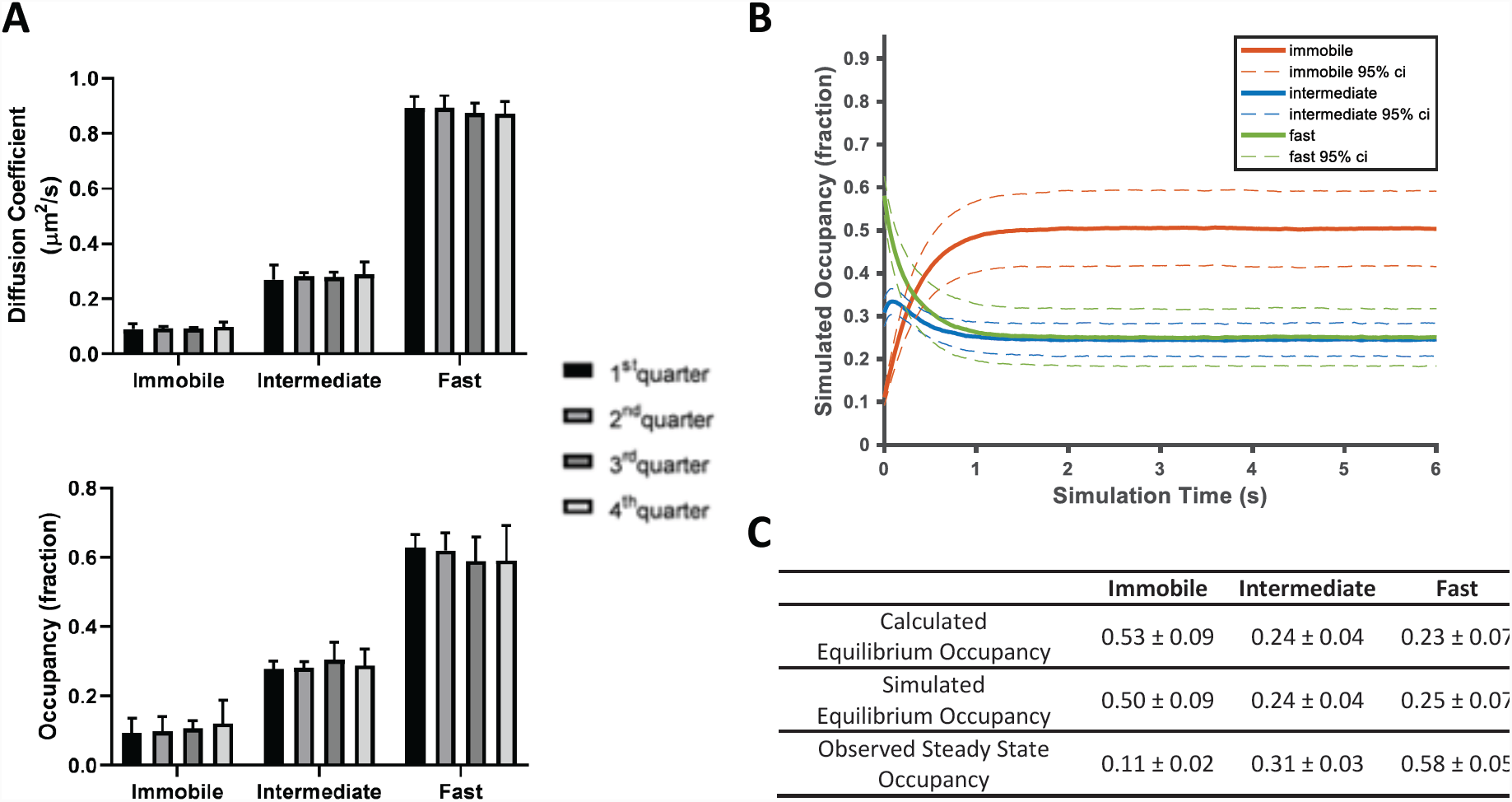
KRas diffusion on the cell membrane is in a non-equilibrium steady state. A) Time invariance of the KRas diffusion model. A single ∼20 min spt-PALM dataset was segmented into four quarters with each quarter containing ∼10,000 trajectories (in ∼5 mins), each analyzed separately using vbSPT to obtain the model parameters such as the diffusion coefficients (upper panel) and the state occupancies (lower panel). Results from multiple spt-PALM datasets were grouped and plotted; B) Temporal evolution of the KRas diffusion model in simulated runs. The system was setup according to the experimental model parameters (number of states, state occupancies, diffusion coefficients, and state transition rates) as shown in Figure 2A. The system was then allowed to evolve based on the input, with the new state occupancies recorded every time step (12 ms) and plotted (see *Methods*). Similar to Figure 2A, only movies with minimum of 30,000 trajectories were simulated; C) Table summarizing the calculated, simulated, and experimentally observed occupancies for each of the states. *All error represents 95% CIs.

To further characterize the NESS, we calculated the mass flow for each of the three KRas diffusion states as the change in state occupancy per time interval. A positive net flow rate or a ratio of in-vs outflux greater than one indicates an accumulation of mass for the state, while a negative flow rate or a ratio of flux less than one indicates the opposite. As shown in Figures 5A & 5B, within the NESS there is a net influx of KRas molecules into the immobile state and a net outflux of molecules out of the fast state, whereas the in- and out-fluxes for the intermediate state are comparable. We also calculated the mass flow for each of the three arms in the diffusion model in Figure 2A – in the clockwise direction, it would be the flow from the fast state to the intermediate state (F to N), intermediate to immobile (N to I), and immobile to fast (I to F). The results of this calculation are shown in Figure 5C, where a positive value in the *y* axis (net mass flow between a pair of states) indicates mass flow in the designated direction and a negative value indicates flow in the opposite direction. Consistent with results in Figures 5A & 5B, the dominant net mass flow through the NESS is unidirectional from the fast state to the intermediate to the immobile state (Fig. 5C), with minimal ‘leakage’ from the fast to the immobile state.

**Figure 5.**
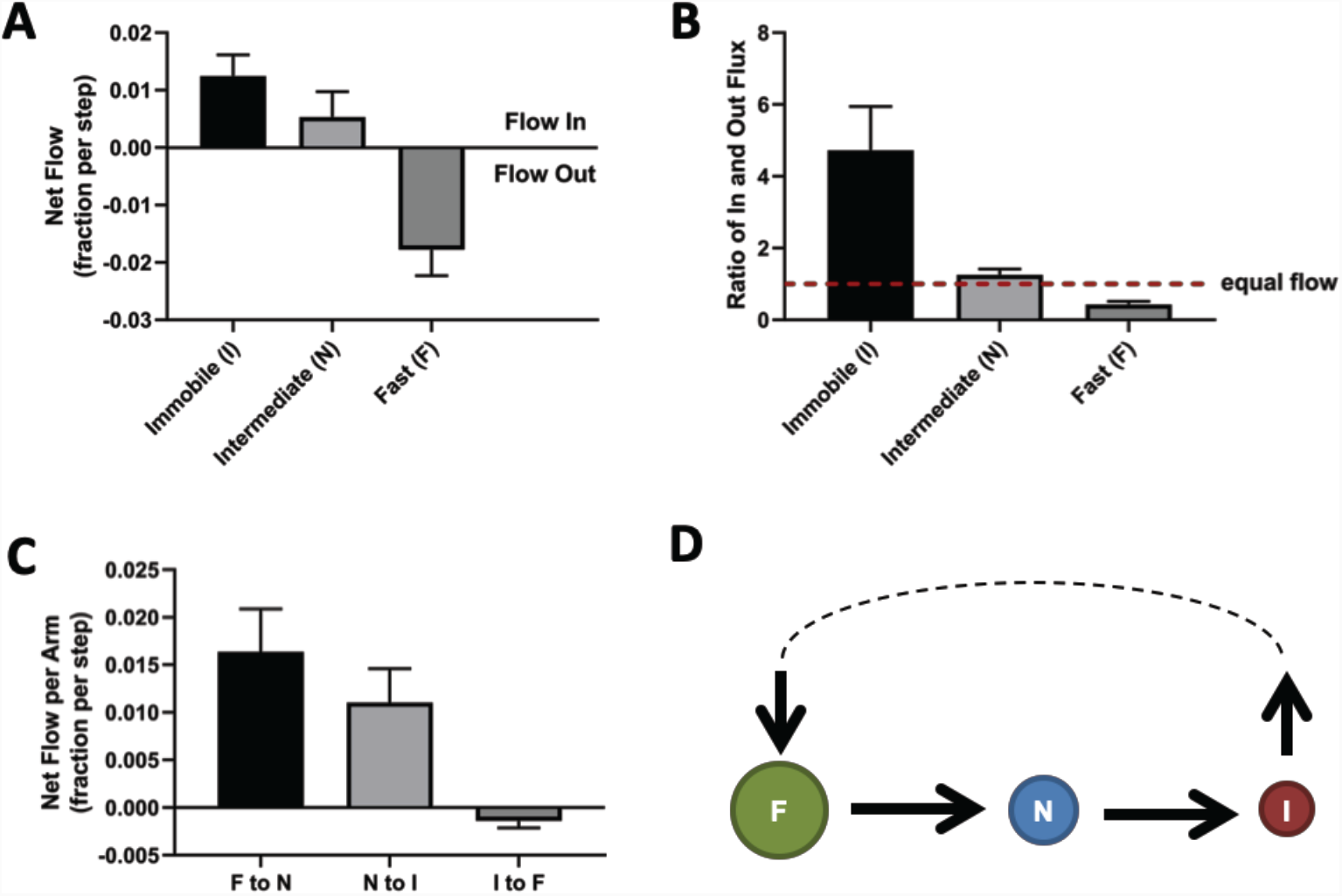
Directional mass flow between the KRas diffusion states. A) Net mass flow per state, defined as the difference between the influx (positive) and the outflux (negative) for each state and expressed as the fraction (of total KRas population) entering (positive, flow in) or leaving (flow out, negative) the state per time interval; B) Ratio of in- and outflux for each state. A ratio of one (dashed line) represents equal in- and outflux for the state, greater than one represents more influx than outflux, and less than one represents net out-flux of mass from the state; C) Net mass flow per arm (pair of states) in the KRas diffusion model (Figure 2A). F to N and N to I are not significantly different. The states were ordered in a clock-wise direction, and the net mass flow in the direction was calculated as the difference between forward and backward mass flows, with a positive value indicating net flow in the indicated direction and a negative value the opposite direction; D) Model for KRas trafficking between the diffusion states and between the membrane system and the environment (cytosol). Arrows indicate the directional mass flow, and the dashed line represents unknown mechanisms connecting the fast and the intermediate states. *All error bars are 95% CIs.

These results are consistent with the simulated relaxation to equilibrium shown in Fig. 4B, where the immobile and the fast diffusion states changed occupancies the most. Therefore, for the KRas NESS system to be sustainable over time as we observed experimentally, KRas would need to be replenished into the fast diffusion state and removed from the immobile state. Indeed, KRas has previously been shown to undergo a constant exchange between the plasma membrane and the cytosol, where internalized KRas is collected at recycling endosomes and transported back to the plasma membrane^50,51^. Our results suggest that the loss of KRas from the membrane would primarily be through the immobile state, and the replenishment through the fast state. At present, it is unclear whether the intermediate state has no exchange with the cytosol or has active exchange with equal gain and loss. Accordingly, the membrane trafficking of KRas should follow the model presented in Fig. 5D, where the arrows indicate the net mass flow between the connected states as well as between the states (F or I) and the environment (cytosol).

### KRas diffusion model is invariant over a range of expression levels

Next, we sought to investigate whether experimental conditions such as expression level would alter the diffusion properties of KRas. An important observation on the nanocluster (multimer) formation of Ras is that the fraction of clustered molecules remains constant over a broad range of expression levels^17^. This unusual property has led to two hypothetical mechanisms of membrane nanocluster formation: one based on protein self-nucleation^17^ and another involving transient actomyosin activity^52^. These energy driven mechanisms are in contrast to passive localization of Ras to pre-existing membrane nanodomains via diffusion, which would result in concentration-dependent multimer formation. To date, the mechanism through which Ras forms multimers remains controversial. Given that membrane nanodomains are associated with the intermediate and the immobile states of KRas, we asked whether the partitioning of KRas in these nanodomains – that is, the fraction of KRas in the corresponding diffusion states – would change with KRas expression level.

To address this question, we induced PAmCherry1-KRas G12D at a range of expression levels using different Dox concentrations (Fig. 1A). Similar to our previous report^12^, the expression level of PAmCherry1-KRas G12D responded well to varying Dox concentrations in the isogenic cells used in this study, with the protein expression at 0 ng/mL being extremely low (only due to occasional leakage in tetR suppression) and that at 10 ng/mL about 5-10 fold higher than endogenous KRas. When measured in terms of protein density at the membrane, the endogenous level of KRas (matched by PAmCherry1-KRas G12D at ∼2 ng/mL Dox) in U2OS cells is ∼60 molecules per μm^2^, and the tuning range corresponds to <10 molecules per μm^2^ at 0 ng/mL Dox to >300 molecules per μm^2^ at 10 ng/mL Dox, or equivalently ∼1/6 to ∼5 times of the endogenous levels of KRas^12^.

By comparing estimated model parameters using spt-PALM data of PAmCherry1-KRas G12D at different Dox concentrations, we found that KRas diffusion properties remained essentially the same across the range of expression levels investigated (Figs. 6A-B and Suppl. Fig. 4). This model invariance is reflected across all conditions: not only was a three-state model optimal for describing the diffusion behavior of KRas as judged with vbSPT (not shown) and with CDF (Suppl. Fig. 9), but the diffusion coefficients of each state, the state occupancies, as well as the transition probabilities between each pair of states, are indistinguishable within the error bars.

**Figure 6.**
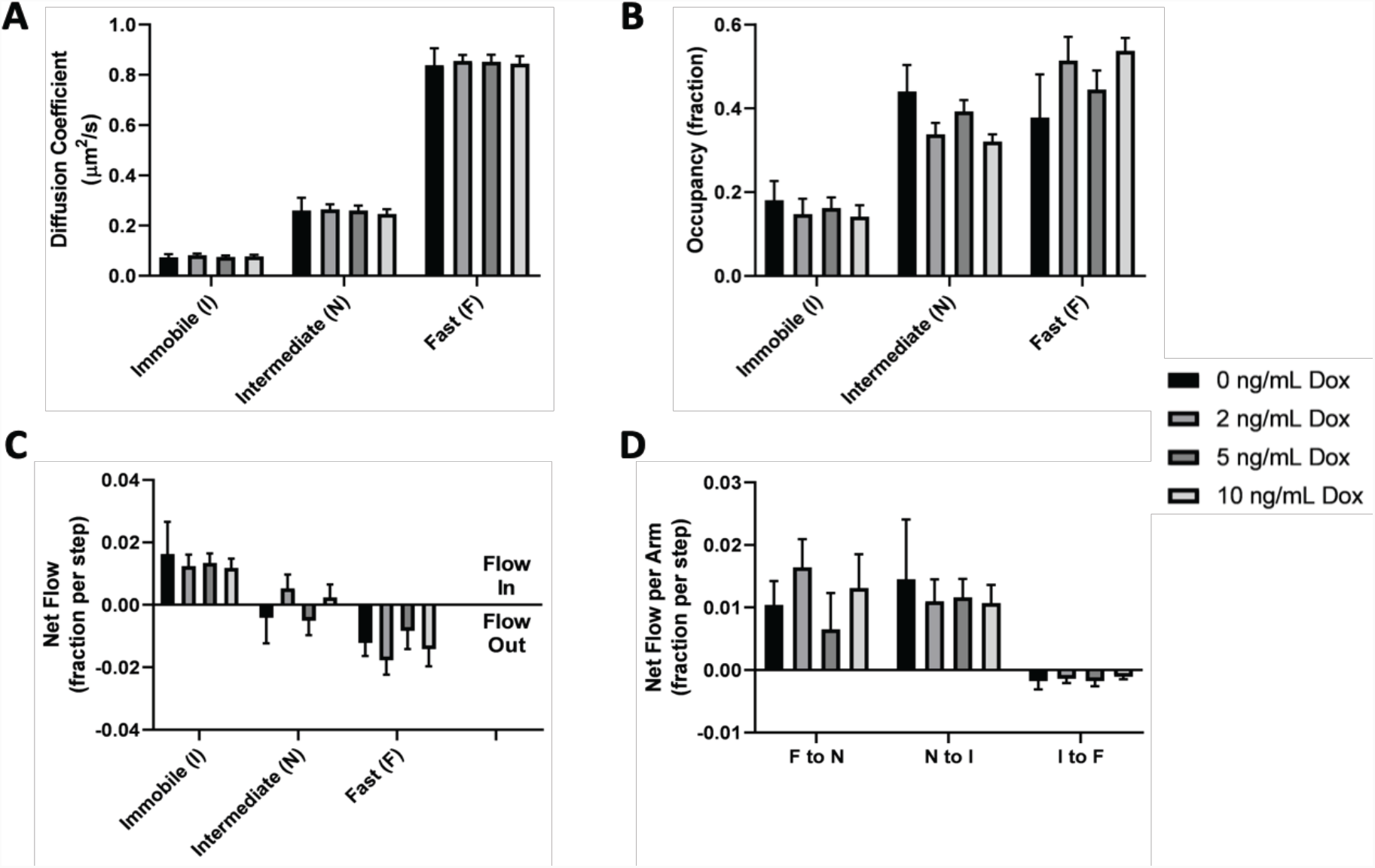
KRas diffusion properties remain constant over a broad range of expression levels. Spt-PALM trajectories of KRas were acquired at 12 ms/frame after inducing the cells at 0, 2, 5, and 10 ng/mL Dox for 36-48 hours, and the diffusion models were inferred as described previously using vbSPT. All aspects of the diffusion model discussed earlier, including diffusion coefficients (A), state occupancies (B), net mass flow per state (C), and net mass flow per arm (pair of states, D) at the different Dox concentrations were analyzed and compared. Error bars represent 95% CIs.

As expected, the net mass flow rates (expressed as the change in state occupancy per time interval) of KRas within the system also remained the same across all the Dox concentrations (Figs. 6C-D). A similar observation was made when we acquired the trajectories at 35 ms/frame (Suppl. Fig. 4D). Thus, we concluded that the KRas diffusion model and the associated trafficking on the membrane remains constant over a broad range of KRas expression levels. Equivalently, the partitioning of KRas in each of the three diffusive states – and the corresponding membrane domains – is stable and independent of its membrane density. This result is consistent with the prior observation that the fraction of Ras in multimers remains constant at widely varying membrane densities^17^.

## Discussion

Membrane nanodomains have been implicated in the regulation of many membrane-resident cellular processes such as Ras signaling^1–6^, but studying the complex and heterogeneous membrane compartments in a living cell has remained a challenge. Using high-throughput SPT and detailed trajectory analysis, we were able to uncover rich details of how KRas localizes and interacts with the membrane. Our results suggest that KRas diffusion on the membrane is best recapitulated with a model that comprises three states – a fast state, an immobile state, and a previously unknown intermediate state. Leveraging the large number of diffusion trajectories, we were able to map the locations where KRas exhibits specific diffusion states. These maps revealed membrane nanodomains corresponding to the intermediate and the immobile states of KRas. The intermediate nanodomains encompass the immobilization sites in a nested configuration, such that KRas almost always transitions between the fast and the immobile states through the intermediate state. We also found that KRas membrane diffusion is in a non-equilibrium steady state, with KRas constitutively removed from the membrane through the immobile sites and replenished as fast diffusing molecules, potentially coupled to KRas trafficking via endocytosis and recycling. Importantly, partitioning of KRas into the three states remains invariant over a wide range of KRas expression levels, demonstrating that KRas diffusion and trafficking through the three mobility states and associated nanodomains is in a maintained, homeostatic condition. Together, these data start to paint a clear picture of the spatiotemporal dynamics of KRas on the membrane, providing the basis for understanding the mechanisms of Ras multimer formation and signaling.

Based on these findings, we propose a new model for Ras membrane diffusion and trafficking as shown in Fig. 7. In this model, Ras experiences at least three types of membrane environments: a ‘regular’ membrane region in which Ras freely diffuses with large step sizes, a ‘transition zone’ or intermediate domain with increased viscous drag and reduced step size, and within the latter an ‘immobilization’ site where Ras interacts with relatively static structures (such as endocytic vesicles). Both the transition zones and the immobilization sites have finite lifetimes, some up to minutes, during which freely diffusing KRas molecules could enter the transition zone, slow down, then either return to the fast state or become trapped in the immobilization sites. During entrapment, a fraction of the trapped KRas molecules leaves the plasma membrane to enter a constitutive cycle of KRas trafficking. This is in agreement with the current understanding that the rate of KRas removal from the membrane through endocytosis is a concentration dependent process, and the localization of KRas at the plasma membrane is an energy driven, PDEδ and Arl2 mediated enrichment of KRas in recycling endosomes which collect and transport KRas back to the plasma membrane^50,51^. Our work adds important details to this trafficking model in that the removal of KRas from the plasma membrane likely occurs during the entrapment phase and its recycling primarily takes place in membrane regions conferring fast mobility. Additionally, the transient entrapment of KRas could also provide an effective mechanism to locally concentrate Ras molecules to facilitate multimer formation, which arguably is a critical step for signaling^22,53,54^. Thus, the various membrane nanodomains directly influence the mobility, trafficking, and potentially multimer formation and signaling of KRas.

**Figure 7.**
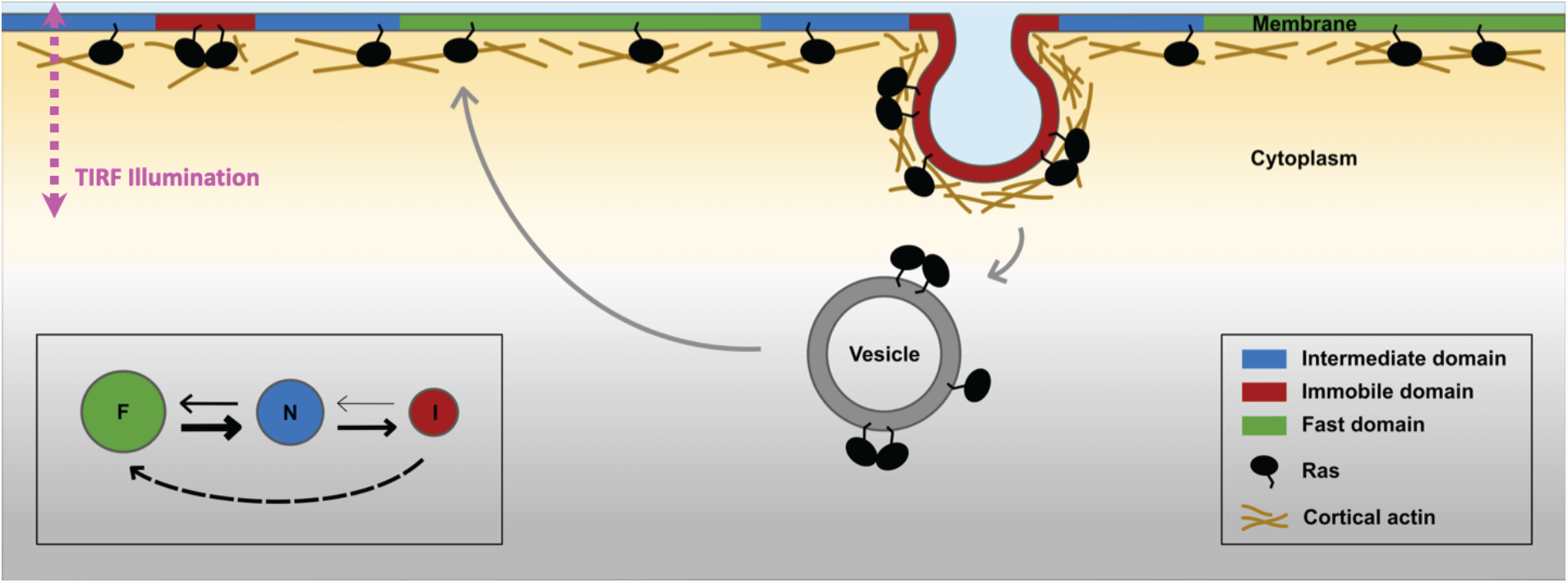
Proposed model for membrane nanodomains regulating KRas mobility and trafficking. For KRas, the cell membrane comprises at least three different compartments conferring each of the three diffusion states of KRas, namely the fast (and free), the intermediate, and the immobile diffusion states, depicted as green, blue, and red regions, respectively. The membrane compartments associated with the immobile and the intermediate states of KRas are nanoscopic membrane structures, and at least a subset of the KRas immobilization structures attributed to endocytic vesicles. KRas is continuously removed from the immobile state, for example through endocytosis, and the internalized KRas molecules are subsequently transported back to the membrane as fast diffusing species through recycling. KRas immobilization domains such as endocytic vesicles could locally enrich KRas molecules to facilitate KRas multimer formation and potentially signaling. The arrows in the legend reflect net flow between each state.

The three-state diffusion model proposed in this study refines existing models of KRas membrane diffusion by introducing a previously unresolved intermediate state, and capturing the role of membrane nanodomains in KRas diffusion. While heterogeneous diffusion properties of KRas and other Ras isoforms have been reported, the prior studies lacked the throughput or spatiotemporal resolutions to determine whether two states, namely a fast diffusion state and an immobile state, are adequate to recapitulate KRas diffusion on the membrane. With the diffusion model defined, we were able to subsequently demonstrate that the intermediate and immobile states of KRas are each associated with a distinct membrane domain. The average sizes of the immobile and the intermediate domains of KRas were found to be ∼70 nm and ∼200 nm, respectively, consistent with previous notion that nanoscopic membrane domains regulate Ras organization on the membrane. We note that, although a three-state model best fits our data, the model could still be an over-simplification. Among other possibilities, both endocytic and non-endocytic mechanisms may contribute to the immobilization of Ras but cannot be distinguished based on diffusion properties since Ras is immobile in both cases. In fact, there are also indications of more than one type of intermediate domains judging from the estimated domain size (Fig. 2F and Suppl. Fig. 6A).

A unique feature of the model in Fig. 7 is that the membrane nanodomains associated with the immobile state of KRas are surrounded by those associated with the intermediate state, creating a nested configuration between the two nanodomains. A plausible scenario is that the structures that trap KRas preferentially form in the membrane regions enriched in certain proteins or lipids and/or more densely packed. In this scenario, KRas would be forced to travel through the intermediate zone to access the immobilization structures, explaining the state transition pathway in Figure 2A. This scenario is also consistent with the observation that the intermediate domains are on average much larger in size than the immobile domains, and that the two nanodomains have similar lifetimes (to the extent of our temporal resolution). In support of this hypothetic scenario, a growing body of literature demonstrates the importance of phosphatidylserine in KRas clustering and activation^18,55–57^. In addition to the KRas tail encoding for phosphatidylserine specificity, a significant fraction of phosphatidylserine display slow motion on the membrane as well^56,58^.

Aside from the steady state partitioning of KRas in the different membrane domains, our data also offered important insight into the membrane dynamics of KRas. We measured a constant flow of KRas from the fast state to the immobile state. Without exchanging KRas with the cytosol, this directional flow would have caused net loss of KRas from the fast state and accumulation in the immobile state as described in Figure 4B-C, yet the experimentally observed state configuration (Fig. 2A) remained stable over time (Fig. 4A). We therefore reasoned that KRas needs to be constantly removed from the immobile state (‘sink’) and replenished via the fast state (‘source’), potentially coupled to membrane trafficking such as endocytosis and recycling^50,51,59–61^. Previous studies have shown that endocytosis is a primary mechanism for KRas removal from the plasma membrane^50^. Thus, our data indicate that at least a subset of the immobile domains could coincide with endocytic vesicles. In support of this, the lifetime of the immobilization domains was estimated to be on the order of 2-5 minutes on average (Fig. 3A-B), which is typical of many endocytic systems^62,63^. The exact mechanism of KRas internalization, however, remains incompletely understood at present.

Remarkably, we found that the spatial partitioning of KRas and more generally the diffusion model were invariant over a broad range of KRas expression levels, which coincides with previous observations where the clustered fraction of KRas or HRas was independent of the protein expression level^17,53^. This corroborates the idea that membrane partitioning of Ras and perhaps many other membrane resident molecules are in an actively maintained, homeostatic condition. This intriguing property of certain membrane proteins^17,64^ has drawn much attention and led to at least two mechanistic models of multimer formation, one based on self-nucleation^17^ and the other driven by actomyosin^52^. Both mechanisms assumed the different states of the protein on the plasma membrane to be in an equilibrium. Our results argue that the mass exchange between the plasma membrane and the cytosol breaks the equilibrium and has to be taken into account in order to accurately model the partitioning behavior of membrane proteins. It is possible that the model presented in Fig. 7 alone or in combination with either self-nucleation or actomyosin activity could better describe the observed model invariance of KRas and potentially other membrane proteins. A clear, mechanistic understanding of this property is important to understand how Ras functions on the membrane, since the Ras multimers have been strongly implicated in signaling. Further experimental and computational work along this line is currently underway.

Last but not the least, a fundamental albeit implicit result from the present study is the importance of experimental parameters in accurately determining the diffusion model, a critical step for in-depth analysis of protein dynamics on the membrane. While there are many different software packages for analyzing spt-PALM trajectories, the importance of controlling the particle density during image acquisition has not previously been recognized to our knowledge. Imaging at a per frame particle density of 0.05-0.1 per µm^2^, which is typical for single-molecule localization microscopy, yielded varying estimated model parameters in our early attempts to track KRas with spt-PALM (Suppl. Text). Using simulations, we found the source of variability to be a small fraction of misconnected trajectories mostly caused by fast moving molecules. In order to minimize the misconnected segments, we kept the density of activated PAmCherry1 in each frame to below 0.03 per µm^2^ at an acquisition rate of 12 ms/frame (Suppl. text and Suppl. Fig. 1). With this precaution, we were able to yield a highly consistent diffusion model from trajectories acquired in different cells and under different conditions. This was critical to defining a previously unresolved state with intermediate mobility (D ∼ 0.3 µm^2^/s) and to all subsequent analyses. We recommend the same precautions to be taken for studies of other membrane molecules.

In summary, our work sheds new light on how complex nanodomains organize on the membrane to dictate Ras diffusion and trafficking. The insights gained from this study offer useful guidance to future experiments that aim at determining the molecular and structural identities of the Ras-associated membrane nanodomains and defining the mechanisms of Ras multimer formation and signaling. The results demonstrate the utility of high-throughput SPT and trajectory analysis in uncovering rich details of the spatiotemporal dynamics of Ras on the membrane, which should be readily applicable to studies of other membrane molecules or processes in cellular compartments.

## Acknowledgements

The authors would like to thank many colleagues for their helpful discussions, including those at OHSU (Drs. Laura Heiser, Molly Kulesz-Martin, Pamela Cassidy, and others at the Department of Dermatology) and those at the Frederick National Laboratory for Cancer Research (Drs. Frank McCormick (also at UCSF), Thomas Turbyville, Dwight Nissley, and others). Research in the Nan lab was supported by startup funds from the Knight Cancer Institute at OHSU, the Damon Runyon Cancer Research Foundation, the M. J. Murdock Charitable Trust, and the Prospect Creek Foundation. XN is also supported by a Cancer Systems Biology Consortium (CSBC) grant (U54 CA209988, PI: Joe Gray) and by the Caner Early Detection Advanced Research (CEDAR) Center of the OHSU Knight Cancer Institute. YL is currently supported by OHSU Center for Spatial Systems Biomedicine (OCSSB). DMZ acknowledges support from the OHSU Center for Spatial Systems Biomedicine and the National Science Foundation, under grant MCB 1715823. We also thank Ms. Julia Shangguan (Nan Lab) for the graphics illustrations, and Dr. Martin Lindén (University of Uppsala) for helpful discussions on vbSPT.

## Author contributions

XN conceived of and designed the project. YL and TH performed the live cell imaging experiments. YL, CP, and TH developed algorithms and software for data analysis. YL, CP, TH, and BM did data analysis. LW and YZ performed biochemical experiments and assisted with imaging. YHC, PJSS, JWG provided guidance on data processing, biochemistry, and cancer biology. DMZ and XN supervised the project.

## Methods

### Cell line

KRas G12D was genetically fused to PAmCherry1, a red fluorescent protein, to ensure high labeling specificity and efficiency. The PAmCherry1-KRas G12D coding sequence is placed under a CMV promoter regulated by the TetOn operon. The construct was transduced via lentivirus into an isogenic U2OS-tetR cell line that constitutively expresses tetR. Single cell clones were subsequently isolated and screened to yield isogenic cell lines that express the PAmCherry1-KRas G12D fusion protein under doxycycline (Dox) regulation.

### Cell treatment for single particle tracking

Cells were grown in fluorobrite DMEM (Thermo Fisher Scientific A1896701) with 10% FBS in eight well Lab-Tek chambers and Dox induced 1.5 days before imaging. Cells were serum starved for at least 12 hours prior to the movie acquisition and did not exceed 24 hours of serum starvation.

### Particle density

We found that particle densities higher than 0.03 µm^-2^ per frame under our experimental conditions (12 ms/frame with the fastest diffusion rate at ∼1 µm^2^/s) led to occasionally misconnected trajectories, and that even a small fraction of such misconnected trajectories could lead to incorrect model outputs with vbSPT (Suppl. Text and Suppl. Fig. 1). In addition, the threshold for maximum displacement between adjacent frames also had an impact on trajectory misconnection, although to a lesser extent for the values tested using simulated trajectories (Suppl. text and Suppl. Figs. 1 & 2). Thus, for diffusion model construction, we chose to use a high frame rate (12 ms/frame) and a low particle density (< 0.03 µm^-2^) to eliminate misconnected trajectory segments while maintaining a sufficient number of trajectories. However, it is beneficial to obtain more trajectories to accurately infer the model parameters with vbSPT, especially for the transition probabilities^36^. As discussed in Suppl. Figure 4, the diffusion coefficients and the occupancies typically converged with only a few thousand trajectories, but the transition probabilities required significantly more trajectories to converge. Thus, we usually acquired spt-PALM data at higher particle densities once the model size has been defined; for these datasets, we safely enforced a three-state model during vbSPT data analysis, since the true diffusion model should not depend on the frame rate or the particle density. This strategy allowed much more flexibility in spt-PALM data acquisition and robustness in the subsequent analyses.

### Trajectory connection for single particle tracking

We constructed single-molecule diffusion trajectories of PAmCherry1-KRas G12D by connecting the centroid positions of the same particles in successive frames. Particles in adjacent frames were deemed to be the same particle if their centroids were within a certain threshold distance. To define the threshold distance, we first constructed the trajectories using a large (∼2,000 nm) distance, from which a step size histogram could be obtained (see Suppl. Fig. 2). The step size histogram from PAmCherry1-KRas G12D typically consists of two segments; signal and noise. The first segment comprises the signal with the first peak around ∼70 nm and extending to ∼500 nm, and all step sizes beyond ∼500 nm was attributed to noise originating from misconnected trajectories generated by the unrealistically large threshold distance. Based on this histogram, we reconstructed the diffusion trajectories using 500 nm as the threshold distance for 12 ms frame acquisition, and 800 nm for 35 ms frame rate movies (using the same method). A new step size histogram was then obtained, which was essentially identical to the first segment of the original step size histogram, confirming that the new threshold distance eliminated most of the misconnected trajectories. The step size histograms of trajectories obtained under the same conditions were also highly consistent, allowing us to set the same threshold value for each condition. Trajectories were terminated if multiple particles were found within the threshold distance in the next frame. Further, all movies acquired at 12 ms frame rate had the additional constraint of having fewer than 0.03 particles/µm^2^ for every frame to lower the chance of misconnecting two different particles in adjacent frames. Thus, all resulting trajectories were constructed without ambiguity. Discussed in further detail in the Suppl. Figs. 1 and 2.

### 2D Markov simulation

We relied on 2D simulations that mimic experimental observations for both experiment optimization and as controls for some of the analysis. Simulations were used to determine the thresholds used for trajectory synthesis (particle density threshold, Suppl. Fig. 1, and connection distance threshold, Suppl. Fig. 2), as well as a negative control to disprove the null hypothesis for spatial clustering (Fig. 2 and 3) and equilibrium analysis (Fig. 4).

The inputs to the simulations were experimentally derived diffusion parameters: number of trajectories, diffusion coefficients, occupancies, transition matrix, frame rate, and the trajectory density. The trajectory density and the number of trajectories are used to determine the width of the simulation space. At the start of the simulation, every particle is randomly assigned a coordinate and a state based on the occupancies. Once a state is assigned, particles are assigned new coordinates by drawing displacements for each dimension from the corresponding *X ∼ N(0, 2Dt)*, where each state has a different diffusion coefficient. At the next time step, a new state is randomly assigned to every particle based on its current state and the transition probability matrix. This process is repeated for the total simulation time. When the simulation was used as the negative control (Figs. 2-4), the simulation was ran for every single movie acquired and the results were compared to the experiment.

### State assignment and averaging

States for each trajectory segment were assigned using vbSPT (contained in field est2.sMaxP, refer to the vbSPT manual). The state assignment is based on trajectory displacements, not the coordinates (e.g. if a trajectory has 3 coordinates, then 2 states are returned for the 2 steps). In order to prevent over counting for the pair correlation analysis (Figs. 2 & 3), in the case of a single molecule staying in the same domain for multiple frames, we averaged all of the coordinates (including both ends) that were assigned the same state for consecutive time points in a single trajectory.

### Pair correlation function

Pair correlation function, or g(r), in general, measures the deviation of the particle density from the expected value from a reference particle as a function of distance. More specifically, g(r) was calculated for each particle by counting the number of other particles within a circular shell at distance of r and r + 10 nm and dividing by the expected number of particles assuming uniform distribution. Therefore, when the observed number of particles for a given distance is equal to the expected number of particles given complete spatial randomness, *g(r)* = 1 and signifies random distribution of particles. Accordingly, *g(r)* > 1 indicates clustering behavior since there are more observed particles around each particle than expected, and *g(r)* < 1 represents cases where there are fewer particles than expected. Every movie was sliced into non-overlapping time segments (1, 5, 10, 20 min) and the average position for each state segment was extracted (as described in State Classification and Averaging) such that every coordinate represented a continuous track for an individual particle in a domain. Therefore, the coordinates used to calculate the pair correlation function represented either different particles that visited the same domain or the same particle that left the domain and returned at a later time. The resulting coordinates were separated into each of the three states, and the *g(r)* was calculated for the coordinates of a given state within the given time slice. In cross pair correlation function analysis, *g(r)* was calculated for a given pair of different states.

## References

1. Grecco, H. E., Schmick, M. & Bastiaens, P. I. H. Signaling from the living plasma membrane. Cell 144, 897–909 (2011).

2. Staubach, S. & Hanisch, F.-G. Lipid rafts: signaling and sorting platforms of cells and their roles in cancer. Expert Rev. Proteomics 8, 263–77 (2011).

3. Simons, K. & Toomre, D. Lipid rafts and signal transduction. Nat. Rev. Mol. Cell Biol. 1, 31–39 (2000).

4. Schmick, M. & Bastiaens, P. I. H. The interdependence of membrane shape and cellular signal processing. Cell 156, 1132–1138 (2014).

5. Mayor, S. & Varma, R. GPI-anchored proteins are organized in submicron domains at the cell surface. Nature 394, 798–801 (1998).

6. Garcia-Parajo, M. F., Cambi, A., Torreno-Pina, J. A., Thompson, N. & Jacobson, K. Nanoclustering as a dominant feature of plasma membrane organization. J. Cell Sci. doi:10.1242/jcs.146340

7. Zhou, Y. & Hancock, J. F. Ras nanoclusters: Versatile lipid-based signaling platforms. Biochim. Biophys. Acta-Mol. Cell Res. 1853, 841–849 (2015).

8. Abankwa, D., Gorfe, A. A., Inder, K. & Hancock, J. F. Ras membrane orientation and nanodomain localization generate isoform diversity. Proc. Natl. Acad. Sci. U. S. A. 107, 1130–5 (2010).

9. Simanshu, D. K., Nissley, D. V & Mccormick, F. RAS Proteins and Their Regulators in Human Disease. Lead. Edge Rev. 170, 17–33 (2017).

10. Cox, A. D. & Der, C. J. Ras history: The saga continues. Small GTPases 1, 2–27 (2010).

11. Güldenhaupt, J. et al. N-Ras forms dimers at POPC membranes. Biophys. J. 103, 1585–93 (2012).

12. Nan, X. et al. Ras-GTP dimers activate the Mitogen-Activated Protein Kinase (MAPK) pathway. Proc. Natl. Acad. Sci. U. S. A. 112, 7996–8001 (2015).

13. Spencer-Smith, R. et al. Inhibition of RAS function through targeting an allosteric regulatory site. Nat. Chem. Biol. 13, 62–68 (2017).

14. Ambrogio, C. et al. KRAS Dimerization Impacts MEK Inhibitor Sensitivity and Oncogenic Activity of Mutant KRAS. Cell 172, 1–12 (2018).

15. Prior, I. a., Muncke, C., Parton, R. G. & Hancock, J. F. Direct visualization of Ras proteins in spatially distinct cell surface microdomains. J. Cell Biol. 160, 165–70 (2003).

16. Inouye, K., Mizutani, S., Koide, H. & Kaziro, Y. Formation of the Ras dimer is essential for Raf-1 activation. J. Biol. Chem. 275, 3737–40 (2000).

17. Plowman, S. J., Muncke, C., Parton, R. G. & Hancock, J. F. H-ras, K-ras, and inner plasma membrane raft proteins operate in nanoclusters with differential dependence on the actin cytoskeleton. Proc. Natl. Acad. Sci. U. S. A. 102, 15500–5 (2005).

18. Zhou, Y. et al. Membrane potential modulates plasma membrane phospholipid dynamics and K-Ras signaling. Science (80-.). 349, 873–876 (2015).

19. Chung, J. K. et al. K-Ras4B Remains Monomeric on Membranes over a Wide Range of Surface Densities and Lipid Compositions. Biophys. J. 114, 137–145 (2018).

20. Belanis, L., Plowman, S. J., Rotblat, B., Hancock, J. F. & Kloog, Y. Galectin-1 Is a Novel Structural Component and a Major Regulator of H-Ras Nanoclusters. Mol. Biol. Cell 19, 1404–1414 (2008).

21. Shalom-Feuerstein, R. et al. K-Ras nanoclustering is subverted by overexpression of the scaffold protein galectin-3. Cancer Res. 68, 6608–6616 (2008).

22. Chen, M., Peters, A., Huang, T. & Nan, X. Ras Dimer Formation as a New Signaling Mechanism and Potential Cancer Therapeutic Target. Mini Rev. Med. Chem. 16, 391–403 (2016).

23. Huang, T., Nickerson, A., Peters, A. & Nan, X. Quantitative fluorescence nanoscopy for cancer biomedicine. in Proceedings of SPIE-The International Society for Optical Engineering 9550, (2015).

24. Schmidt, T., Schütz, G. J., Baumgartner, W., Gruber, H. J. & Schindler, H. Imaging of single molecule diffusion. Proc. Natl. Acad. Sci. U. S. A. 93, 2926–9 (1996).

25. Kusumi, A. et al. Paradigm Shift of the Plasma Membrane Concept from the Two-Dimensional Continuum Fluid to the Partitioned Fluid: High-Speed Single-Molecule Tracking of Membrane Molecules. Annu. Rev. Biophys. Biomol. Struct. 34, 351–378 (2005).

26. Saxton, M. J. & Jacobson, K. Single-particle tracking: applications to membrane dynamics. Annu. Rev. Biophys. Biomol. Struct. 26, 373–399 (1997).

27. Lommerse, P. H. M. et al. Single-molecule diffusion reveals similar mobility for the Lck, H-ras, and K-ras membrane anchors. Biophys. J. 91, 1090–1097 (2006).

28. Murakoshi, H. et al. Single-molecule imaging analysis of Ras activation in living cells. Proc. Natl. Acad. Sci. U. S. A. 101, 7317–22 (2004).

29. Lommerse, P. H. M. et al. Single-Molecule Imaging of the H-Ras Membrane-Anchor Reveals Domains in the Cytoplasmic Leaflet of the Cell Membrane. Biophys. J. 86, 609–616 (2004).

30. Manley, S. et al. High-density mapping of single-molecule trajectories with photoactivated localization microscopy. Nat Meth 5, 155–157 (2008).

31. Benke, A., Olivier, N., Gunzenhäuser, J. & Manley, S. Multicolor single molecule tracking of stochastically active synthetic dyes. Nano Lett. 12, 2619–24 (2012).

32. Huang, T. et al. Simultaneous Multicolor Single-Molecule Tracking with Single-Laser Excitation via Spectral Imaging. Biophys. J. 114, 301–310 (2018).

33. Basu, S. et al. FRET-enhanced photostability allows improved single-molecule tracking of proteins and protein complexes in live mammalian cells. Nat. Commun. 9, (2018).

34. English, B. P. & Singer, R. H. A three-camera imaging microscope for high-speed single-molecule tracking and super-resolution imaging in living cells. Proc. SPIE-the Int. Soc. Opt. Eng. 9550, 955008 (2015).

35. Cutler, P. J. et al. Multi-color quantum dot tracking using a high-speed hyperspectral line-scanning microscope. PLoS One 8, e64320 (2013).

36. Persson, F., Lindén, M., Unoson, C. & Elf, J. Extracting intracellular diffusive states and transition rates from single-molecule tracking data. Nat. Methods 10, 265–9 (2013).

37. Hansen, A. S. et al. Robust model-based analysis of single-particle tracking experiments with Spot-On. Elife 7, 1–33 (2018).

38. Liu, Z., Lavis, L. D. & Betzig, E. Imaging Live-Cell Dynamics and Structure at the Single-Molecule Level. Mol. Cell 58, 644–659 (2015).

39. Jaqaman, K. et al. Robust single-particle tracking in live-cell time-lapse sequences. Nat. Methods 5, 695–702 (2008).

40. Chenouard, N. et al. Objective comparison of particle tracking methods. Nat. Methods 11, 281– 289 (2014).

41. Shen, H. et al. Single Particle Tracking: From Theory to Biophysical Applications. Chem. Rev. 117, 7331–7376 (2017).

42. Monnier, N. et al. Inferring transient particle transport dynamics in live cells. Nat. Methods 12, 1– 7 (2015).

43. Ito, Y., Sakata-Sogawa, K. & Tokunaga, M. Multi-color single-molecule tracking and subtrajectory analysis for quantification of spatiotemporal dynamics and kinetics upon T cell activation. Sci. Rep. 7, 6994 (2017).

44. Lindén, M. & Elf, J. Variational Algorithms for Analyzing Noisy Multistate Diffusion Trajectories. Biophys. J. 276–282 (2018). doi:10.1016/j.bpj.2018.05.027

45. Newby, J. M., Schaefer, A. M., Lee, P. T., Forest, M. G. & Lai, S. K. Deep neural networks automate detection for tracking of submicron scale particles in 2D and 3D. Proc. Natl. Acad. Sci. USA 115, 9026–9031 (2018).

46. Manzo, C. & Garcia-Parajo, M. F. A review of progress in single particle tracking: from methods to biophysical insights. Rep. Prog. Phys. 78, 124601 (2015).

47. Subach, F. V et al. Photoactivatable mCherry for high-resolution two-color fluorescence microscopy. Nat. Methods 6, 153–9 (2009).

48. Schuitz, G. J., Schindler, H. & Schmidt, T. Single-Molecule Microscopy on Model Membranes Reveals Anomalous Diffusion. Biophys. J. 73, 1073–1080 (1997).

49. Zuckerman, D. Statistical Physics of Biomolecules: An Introduction. (CRC Press, 2010).

50. Schmick, M. et al. KRas localizes to the plasma membrane by spatial cycles of solubilization, trapping and vesicular transport. Cell 157, 459–471 (2014).

51. Schmick, M., Kraemer, A. & Bastiaens, P. I. H. Ras moves to stay in place. Trends Cell Biol. 25, 190–7 (2015).

52. Gowrishankar, K. et al. Active remodeling of cortical actin regulates spatiotemporal organization of cell surface molecules. Cell 149, 1353–1367 (2012).

53. Tian, T. et al. Plasma membrane nanoswitches generate high-fidelity Ras signal transduction. Nat. Cell Biol. 9, 905–14 (2007).

54. Nussinov, R., Tsai, C.-J. & Jang, H. Is Nanoclustering essential for all oncogenic KRas pathways? Can it explain why wild-type KRas can inhibit its oncogenic variant? Semin. Cancer Biol. 0–1 (2018). doi:10.1016/j.semcancer.2018.01.002

55. Cho, K.-J. et al. Inhibition of Acid Sphingomyelinase Depletes Cellular Phosphatidylserine and Mislocalizes K-Ras from the Plasma Membrane. Mol. Cell. Biol. 36, 363–74 (2016).

56. Zhou, Y. et al. Lipid-Sorting Specificity Encoded in K-Ras Membrane Anchor Regulates Signal Output. Cell 168, 239–251.e16 (2017).

57. Ariotti, N. et al. Caveolae regulate the nanoscale organization of the plasma membrane to remotely control Ras signaling. J. Cell Biol. 204, 777–792 (2014).

58. Kay, J. G., Koivusalo, M., Ma, X., Wohland, T. & Grinstein, S. Phosphatidylserine dynamics in cellular membranes. Mol. Biol. Cell 23, 2198–212 (2012).

59. Roy, S., Wyse, B. & Hancock, J. F. H-Ras signaling and K-Ras signaling are differentially dependent on endocytosis. Mol. Cell. Biol. 22, 5128–5140 (2002).

60. Rocks, O. et al. An acylation cycle regulates localization and activity of palmitoylated Ras isoforms. Science 307, 1746–52 (2005).

61. Lu, A. et al. A clathrin-dependent pathway leads to KRas signaling on late endosomes en route to lysosomes. J. Cell Biol. 184, 863–879 (2009).

62. Mayor, S., Parton, R. G. & Donaldson, J. G. Clathrin-independent pathways of endocytosis. Cold Spring Harb. Perspect. Biol. 6, 1–20 (2014).

63. Kaksonen, M., Toret, C. P. & Drubin, D. G. A modular design for the clathrin- and actin-mediated endocytosis machinery. Cell 123, 305–320 (2005).

64. Sharma, P. et al. Nanoscale organization of multiple GPI-anchored proteins in living cell membranes. Cell 116, 577–89 (2004).

