## Supplementary Information for "High-throughput single-particle tracking reveals nested membrane nanodomain organization that dictates Ras diffusion and trafficking"

#### List of supplementary material

1. **Supplementary Text & Figure 1.** Impact of particle density on diffusion model reconstruction.
2. **Supplementary Figure 2.** Impact of the connection distance threshold on vbSPT model output.
3. **Supplementary Figure 3.** vbSPT model output on experimental spt-PALM datasets acquired at high particle densities.
4. **Supplementary Figure 4.** Impact of frame rate and total number of trajectories on vbSPT model output.
5. **Supplementary Figure 5.** Photon yield and localization accuracy at the different frame rates used in this work.
6. **Supplementary Figure 6.** Spatial analysis of KRas membrane domain properties using data acquired at 35 ms per frame.
7. **Supplementary Figure 7.** Temporal evolution of the membrane domains associated with each KRas diffusive state.
8. **Supplementary Figure 8.** Validating vbSPT output accuracy on simulated trajectories using different model parameter inputs.
9. **Supplementary Figure 9.** The three-state model remains optimal for KRas membrane diffusion over a broad range of expression levels.

10. **Supplementary video 1.** A short 1000 frame clip of a raw single particle tracking experiment of KRas G12D fused to PAmCherry1 acquired at 35 ms frame interval. The mutant KRas was induced with 5 ng/mL doxycycline and expressed in the U2OS cell line.
11. **Supplementary video 2**. A movie of the domain map as described in SI figure 6.

**Supplementary text**

This section addresses how the particle density in each image frame impacts the diffusion model reconstruction. Our investigation into this issue was first prompted by the observation that when using vbSPT to infer the diffusion model based on the experimental spt-PALM data, the model outputs varied in terms of both the model size (i.e., number of diffusive states) as well as the model parameters (e.g. diffusion coefficients), even for data acquired from different cells under the same conditions. Moreover, vbSPT also yielded different models depending on the threshold of step size used for connecting the particle coordinates in adjacent frames into trajectories.

Using simulated trajectories (see *Methods*), we found that the number of particles per raw image frame has a big impact on the diffusion model outputs from vbSPT and CDF. At high particle densities, two different particles from successive frames may be connected leading to some misconnected trajectories and a significant increase of nonexistent, fast-moving particles, which vbSPT would assign to artificial states. As demonstrated in Supplementary Figure 1, misconnection occurs only in a tiny fraction of the trajectories (corresponding to the positive end of the step size histogram) and predominantly skews the fast diffusive states of the model where most ambiguities in trajectory connection arise. This manifest as altered diffusion parameters of the fast states or appearance of non-existent 'fast' states with suspiciously low occupancies.

The severity of trajectory misconnection depends on how fast the particles diffuse, how sparse the particles are in each frame, and how quickly the image acquisition takes place. For a given frame rate, the fastest diffusion coefficient dictates the maximum travel distance of a particle, which in turn affects the step size (particle search distance for trajectory connection) threshold. A large search distance threshold results in an increased likelihood of connecting two unrelated particles, especially when the particle density is high. Hence, for accurate model reconstruction, it is better to use a high frame rate with a small number of particles per frame. This effectively eliminates misconnected trajectories and still yields a sufficient number of diffusion trajectories to accurately determine the model (Suppl. Fig. 4).

We determined the optimal particle density per frame for tracking KRas G12D by testing the performance of our analysis workflow with simulated trajectories at varying particle densities (see *Methods*). We ran 2D simulations of diffusing particles with 2 diffusion states (0.1 and 1 $\mu m^2/s$, referred to as the 'slow' and the 'fast' states, respectively) at 12 ms frame acquisition time and particle densities at 0.03, 0.04, and 0.05 per $\mu m^2$ per frame. Here, the fast diffusion coefficient at 1 $\mu m^2/s$ was used empirically based on previous reports as well as model outputs from existing datasets[1–3]. Particle coordinates generated in the simulations were connected in the same way as the experimental datasets to intentionally introduce errors in trajectory connection (see *Methods*). As shown in Supplementary Figure 1, while the step size histograms at the three different particle densities seemed virtually identical (Suppl. Fig. 1A), vbSPT reported different models for each condition (Supplementary Figures 1B-D), with only the model output at 0.03 per $\mu m^2$ per frame being accurate because it correctly predicts a 2-state model. This suggests that the fraction of misconnected trajectories at the higher particle densities could be too small to be detected by visual inspection but it does lead to incorrect model reconstruction. Starting at 0.04 particles per $\mu m^2$ per frame, vbSPT detected a third state with a diffusion coefficient greater than 1 $\mu m^2/s$ but with virtually null occupancies (Suppl. Figs. 1C&D). Thus, we set the maximum particle density threshold for all of our movies to be 0.03 particles/$\mu m^2$. Interestingly, CDF fitting correctly returned a 2 state model for all the particle densities tested (Suppl. Figures 1E-G), making CDF fitting a useful parallel approach to confirming the model output of vbSPT.

We note that once the diffusion model under a given biological condition is defined by using high frame rate and low particle density data, further analyses could be done using data obtained at higher particle densities and/or lower frame rates (i.e., 'non-ideal' conditions) based on the same model as long as the biological setting remains constant. The diffusion model should be independent of image acquisition speed and the number of particles appearing in each frame and having more particles and/or trajectories typically offers better statistical power for the analyses.

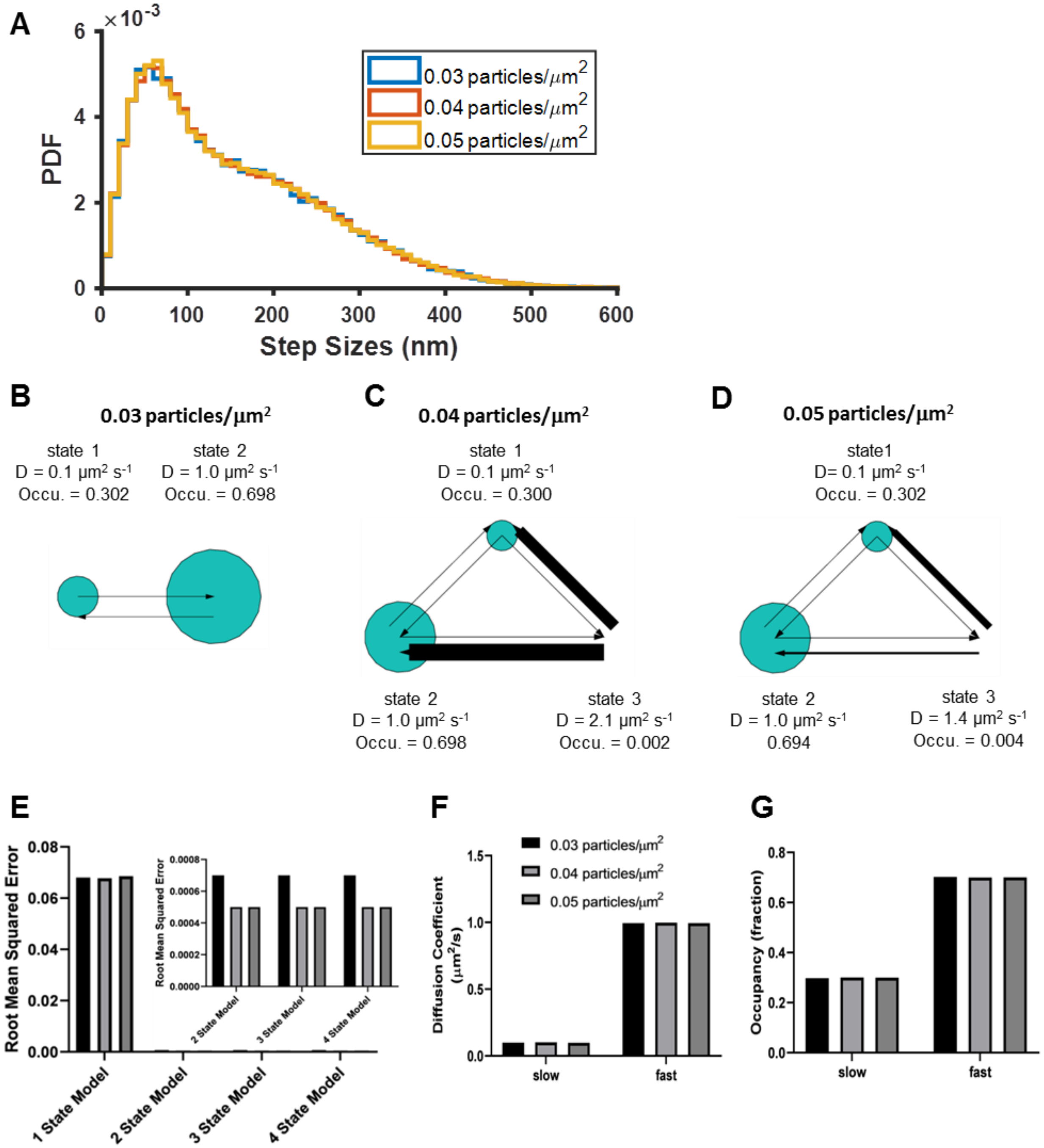


**Supplementary Figure 1. Impact of particle density on diffusion model reconstruction.** Test data were generated by simulating diffusion trajectories of two separate populations of particles with no transitions exhibiting diffusion coefficients of 0.1 and 1 $\mu m^2 s^{-1}$, and occupancies of 0.3 and 0.7, respectively (see *Methods*). About 6-10k trajectories were synthesized (depending on the particle density) with connection distance threshold of 600 nm and analyzed using vbSPT (B-D) or CDF (E-G). A) Histograms of step sizes at 0.03, 0.04, and 0.05 particles/$\mu m^2$ per frame; B-D) show the vbSPT outputs on simulated trajectories at 0.03, 0.04, and 0. 05 particles/$\mu m^2$, returning 2, 3, and 3 state models respectively, with the model parameters displayed next to each state; E) Goodness of CDF fitting at different model sizes, as well as the diffusion coefficients (F) and state occupancies (G) obtained from fitting to a 2-state model.

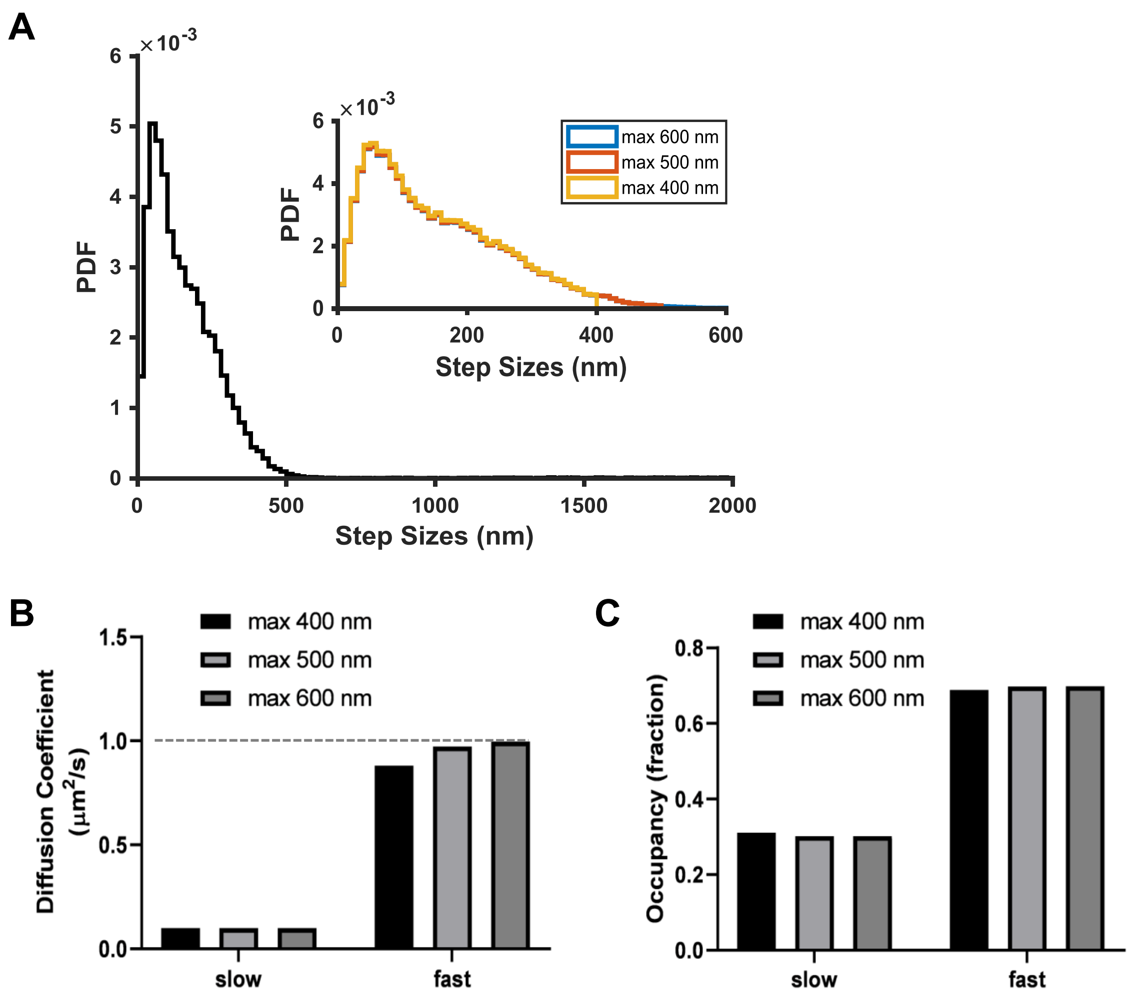


**Supplementary Figure 2. Impact of the connection distance threshold on vbSPT model output.** The maximum connection distance threshold specifies the search radius around each particle in the current frame for its possible locations in the next frame. This is a critical parameter for linking particle coordinates into trajectories but is initially unknown. To address this challenge, we used simulated trajectories at 0.03 particles/$\mu m^2$ per frame as in Supplementary Figure 1, which comprises a two state system with diffusion coefficients at 0.1 and 1 $\mu m^2$/s. We first synthesized trajectories using an unrealistically large connection distance threshold of 2000 nm, and examined the step size distribution of the resulting trajectories (A, main panel), where it became clear that the vast majority of the molecules moved less than 500 nm between frames. Based on this, we chose 400 nm, 500 nm, and 600 nm as the connection distance thresholds and resynthesized the trajectories (A, inset); the histograms essentially overlap except at the large step sizes (blue: 600 nm threshold; red: 500 nm threshold; and yellow: 400 nm threshold). B-C) Comparison of the vbSPT model outputs on trajectories synthesized using the three connection distance threshold values shows that vbSPT was able to pick the correct model size (of 2) at all three settings. However, setting the threshold value at 400 or even 500 nm caused a noticeable truncation in the step size histogram (as shown in A) and resulted in lower diffusion coefficients for the fast state while the 600 nm threshold returned the correct diffusion coefficient; the slow state was not affected. Interestingly, the threshold setting had minimal impact on the resulting outputs for state occupancies (C). These settings were used to guide the trajectory synthesis based on the experimental spt-PALM data.

**A**

| Model Size | Count of Movies with Model Size |
|---|---|
| **4** | 3 |
| **5** | 3 |
| **6** | 8 |
| **8** | 1 |

**B**

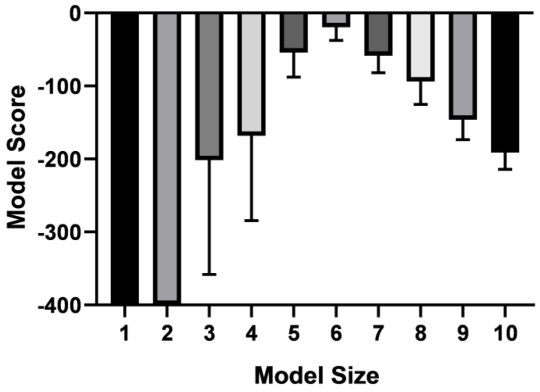


**C**

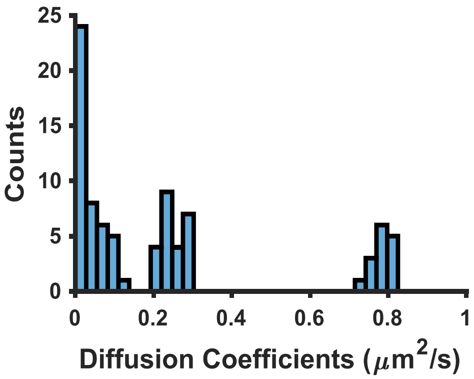


**Supplementary Figure 3. vbSPT model output on experimental spt-PALM datasets acquired at high particle densities.** When spt-PALM datasets of PAmCherry1-KRas G12D in U2OS cells were acquired at particle densities higher than 0.03 per $\mu m^2$ per frame (typically around 0.05 – 0.1 per $\mu m^2$ per frame), vbSPT outputs diffusion models of varying sizes, many reaching 6 or more states (A, B). However, a histogram of the diffusion coefficients of all detected states shows 3 clusters, indicating that a three-state model is still likely the best to recapitulate KRas diffusion (C). Note that the three clusters are centered at diffusion coefficient values similar to those obtained with vbSPT or CDF analysis of spt-PALM datasets acquired at low particle densities (<0.03 particles per $\mu m^2$ per frame) as in Figure 1F, and are especially close to the diffusion coefficients acquired at 35 ms frame rate shown in Supplementary Figure 4A. All data were taken at 35 ms/frame.

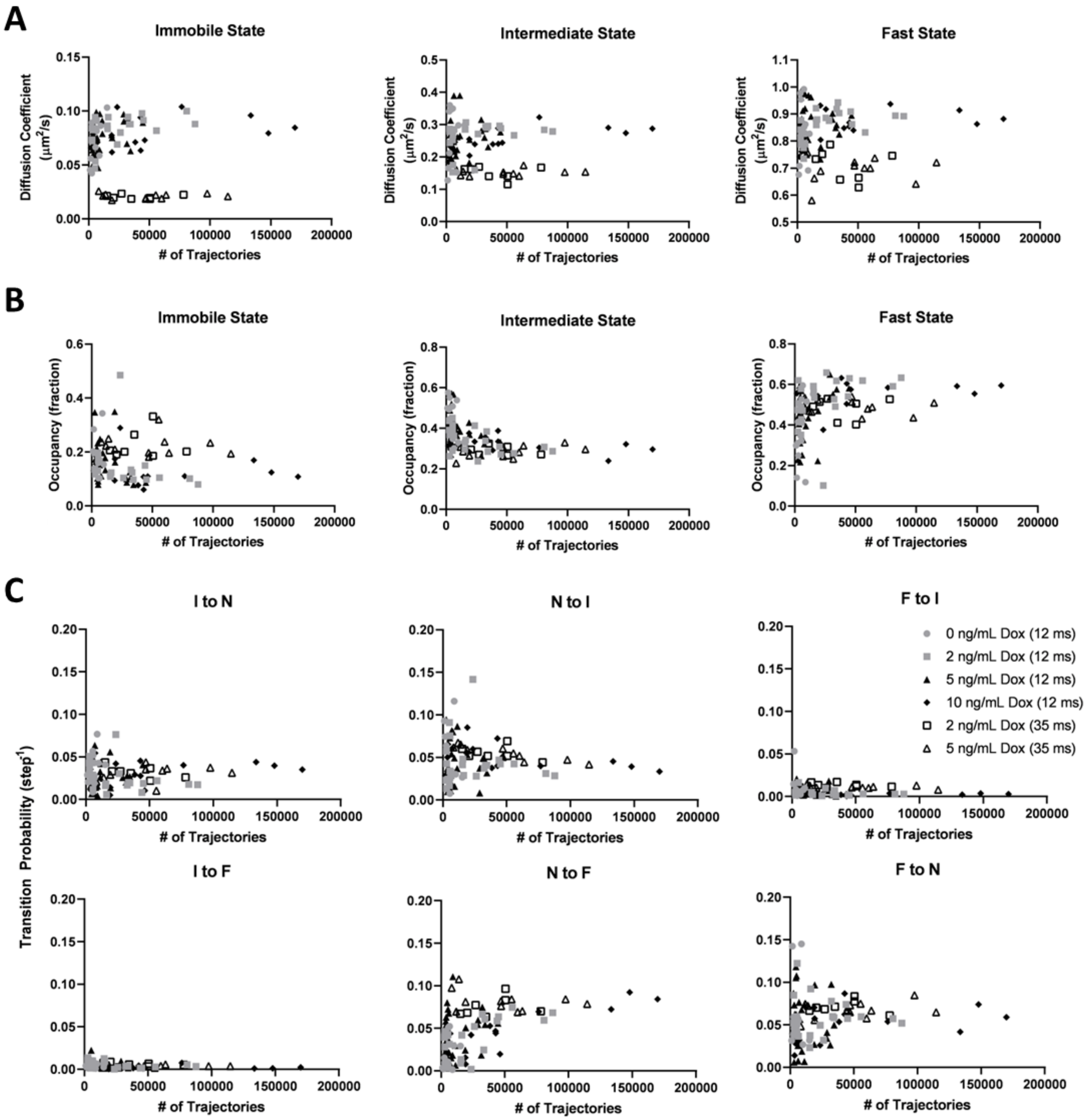

A
Immobile State
Intermediate State
Fast State
Diffusion Coefficient (μm²/s)
### of Trajectories
B
Immobile State
Intermediate State
Fast State
Occupancy (fraction)
### of Trajectories
C
I to N
N to I
F to I
I to F
N to F
F to N
Transition Probability (step⁻¹)
### of Trajectories
0 ng/mL Dox (12 ms)
2 ng/mL Dox (12 ms)
5 ng/mL Dox (12 ms)
10 ng/mL Dox (12 ms)
2 ng/mL Dox (35 ms)
5 ng/mL Dox (35 ms)

D

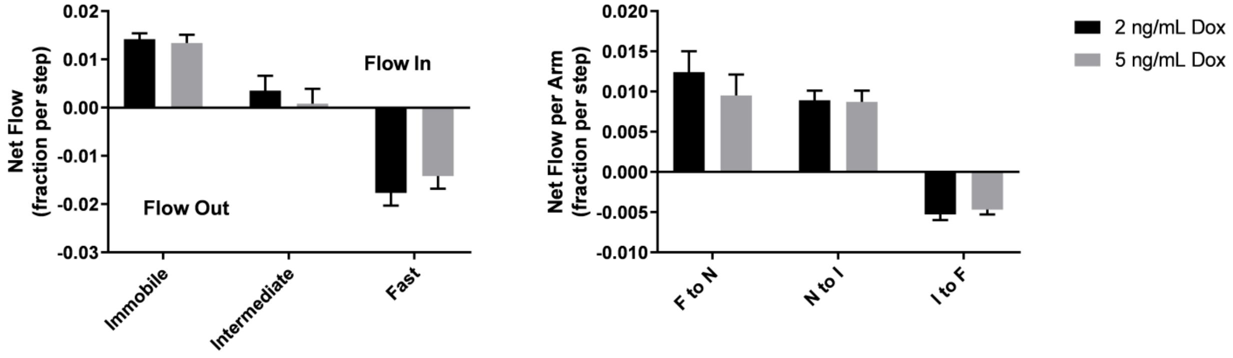


**Supplementary Figure 4. Impact of frame rate and total number of trajectories on vbSPT model output.** All experimental spt-PALM data on PAmCherry1-KRas (in U2OS cells) acquired with 12 or 35 ms frame acquisition times and under 0, 2, 5, or 10 ng/mL Dox concentrations were pooled (symbols as indicated), and vbSPT outputs of the diffusion coefficients (A), state occupancies (B), and state transition probabilities (C) were plotted against the total number of trajectories. As shown in (A) and (B), the diffusion coefficients and the occupancies typically converge relatively quickly at a few thousand trajectories. Additionally, the diffusion coefficients derived from datasets obtained at 35 ms/frame are consistently lower than those obtained with 12 ms/frame datasets, a result of both localization precision (particularly for the immobile state) and trajectory smearing (predominantly for the faster diffusive states). Transition probabilities (C) required more trajectories to converge. However, all model parameters converged at similar values regardless of Dox concentration (i.e., KRas expression level). D) Net flow analysis on the 35 ms/frame dataset for 2 and 5 ng/mL Dox.

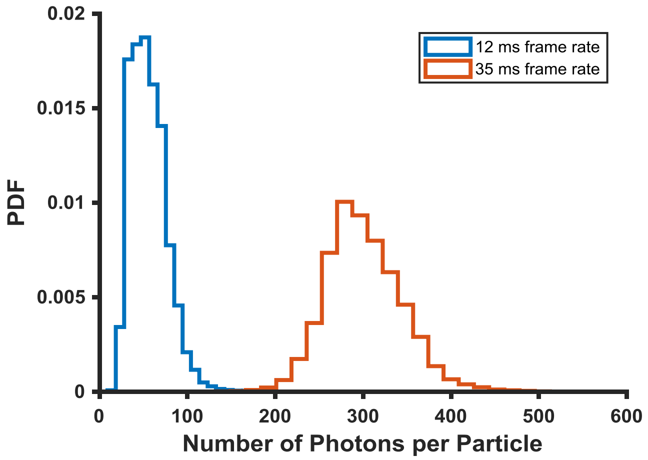


**Supplementary Figure 5. Photon yield and localization accuracy at the different frame rates used in this work.** Photon yields were calculated based on the integrated intensity above background across a 9×9 pixel area for each single-molecule image; the pixel intensity units were converted to the number of photons using hardware specific gain conversion factors. On average, the photon yield for single PAmCherry1 molecules at 12 ms and 35 ms frame acquisition time was ~56 photons and ~301 photons, corresponding to ~26 nm and ~10 nm localization precisions, respectively.

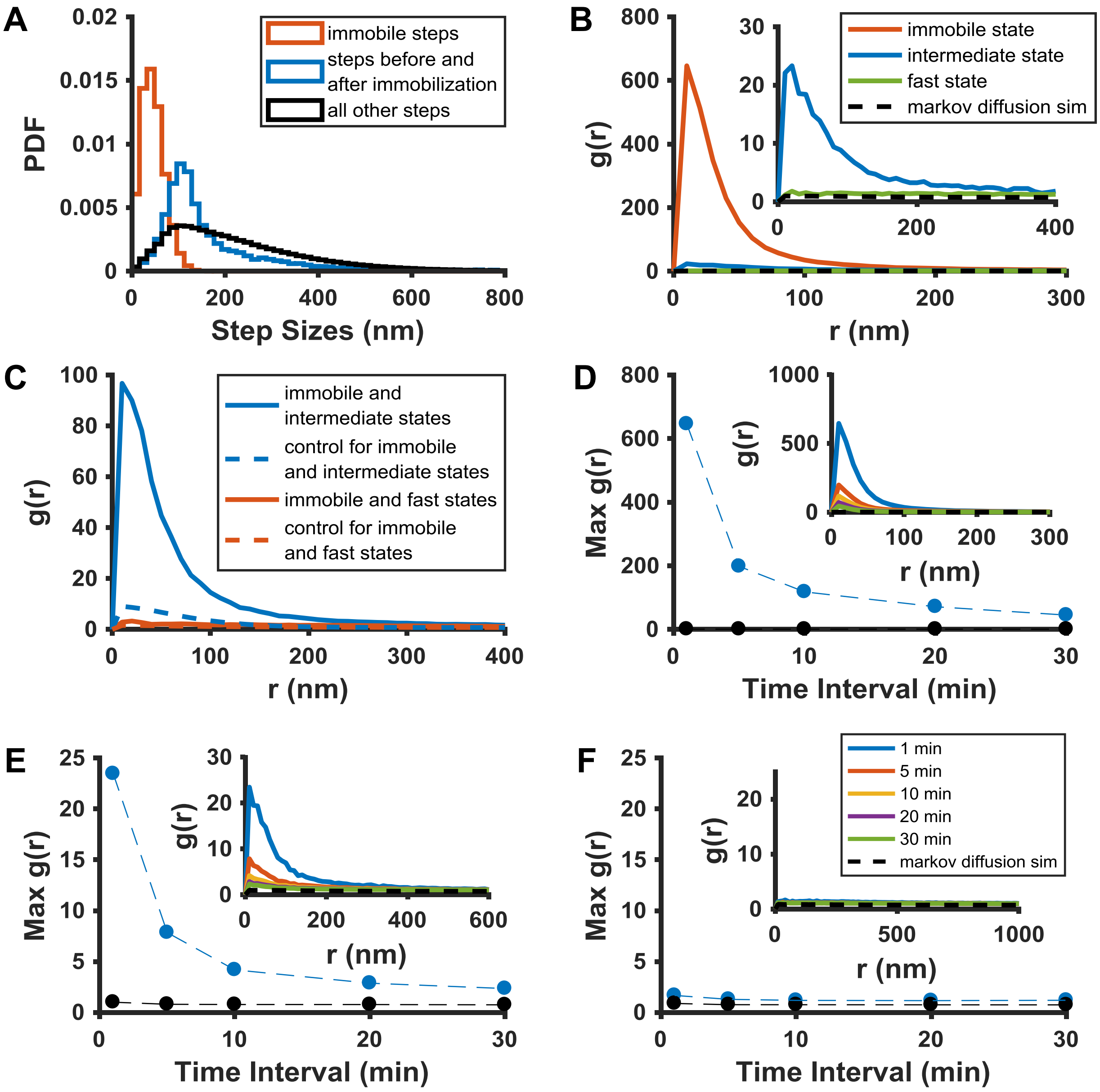


**Supplementary Figure 6. Spatial analysis of KRas membrane domain properties using data acquired at 35 ms per frame.** As spt-PALM data acquired at 35 ms/frame showed better single-molecule localization accuracy than those at 12 ms/frame, we aimed to perform similar analysis of the domain properties to that shown in Figures 2 & 3 using data taken at 35 ms/frame. A) Step size histograms for the immobilization events (red), the steps directly before and after the immobilization events (blue), and all other steps (black); B) Pair correlation analysis on the averaged positions of the three states for one-minute temporal slices of the raw spt-PALM image stack (see *Methods*), shows the same trend as observed with data taken at 12 ms/frame acquisition rate. Note the somewhat reduced spatial correlation for the intermediate domain (state) compared with that obtained with data taken at 12 ms/frame (Figure 2D); C) Cross-correlation analysis between the three membrane domains, performed on the same one-minute slices of the raw spt-PALM image stack; D-F) Peak amplitudes of autocorrelation *g(r)* at different time intervals. The steadily decreasing *g(r)* with increasing time intervals indicates the lifetimes of the immobile and the intermediate domains to be on the order of minutes.

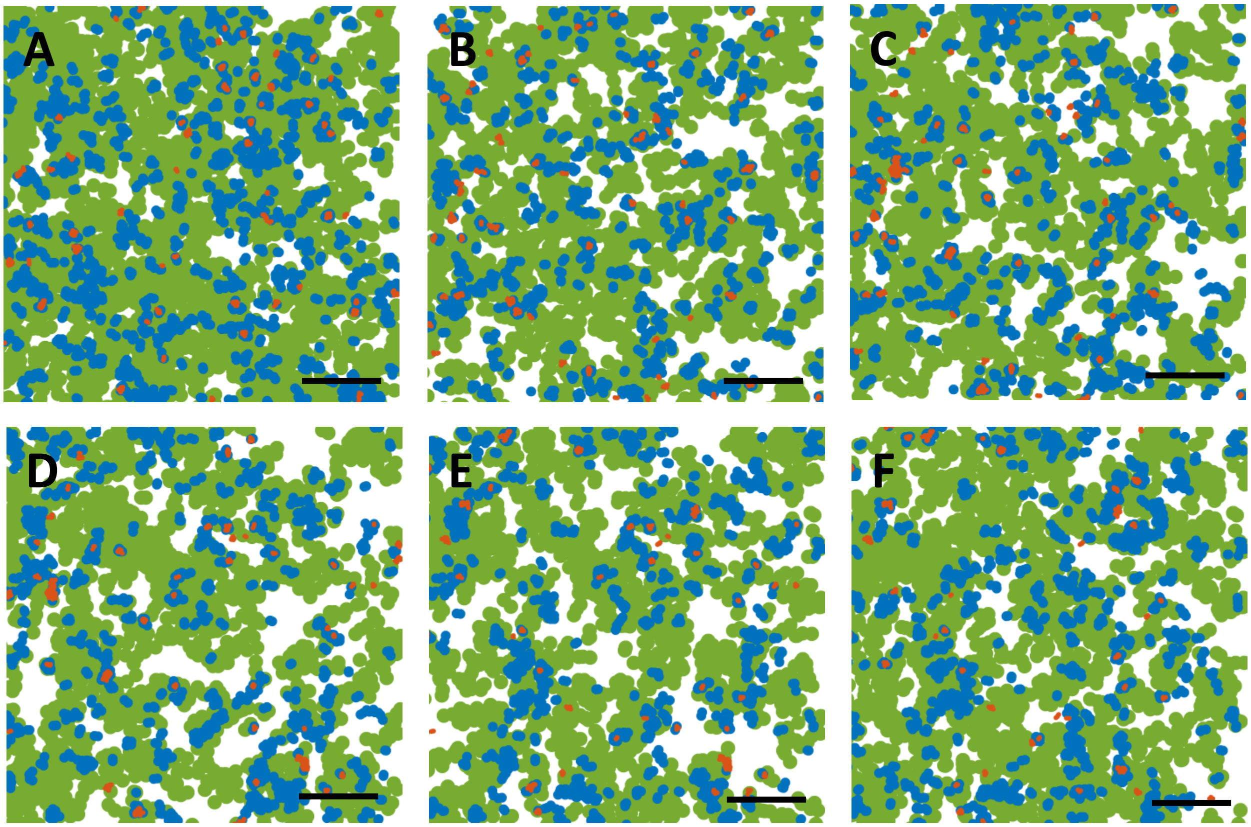


**Supplementary Figure 7. Temporal evolution of the membrane domains associated with each KRas diffusive state.** The three membrane domains associated with the immobile, intermediate, and fast states of KRas are labeled with red, blue, and green, respectively. The domain maps were generated using the same approach as described for Figure 2C (12 ms frame interval), with each panel representing the domain map within a 1 min duration with 0.5 min overlap. Thus, A-C represent total of 3.5 min time period. Of note, the maps were generated without position averaging, and therefore each trajectory contributes 2 or more points (including the beginning and the end) in the corresponding plot. Scale bars, 2 µm. See also supplementary video 2.

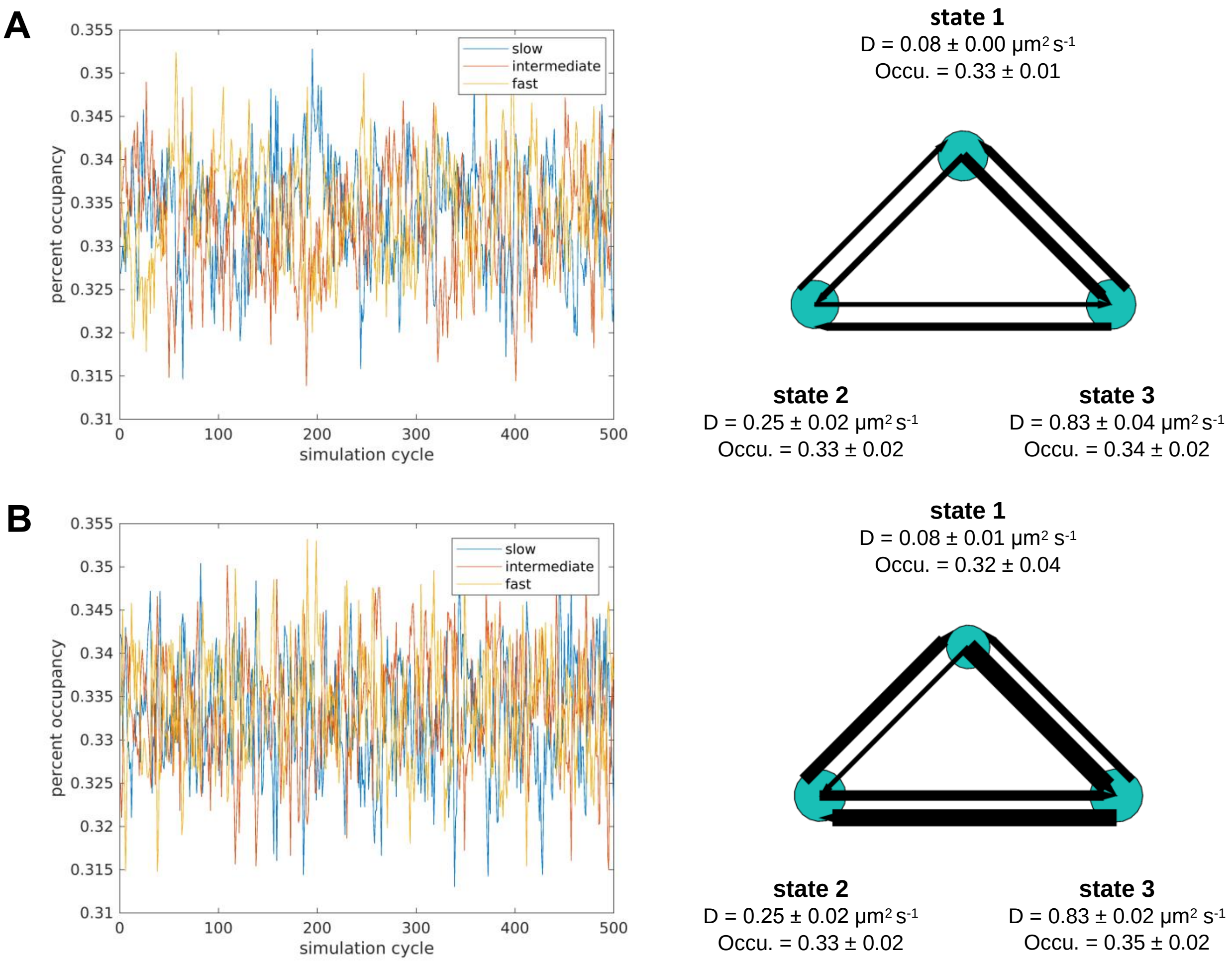


**Supplementary Figure 8. Validating vbSPT output accuracy on simulated trajectories using different model parameter inputs.** We simulated steady state systems using three states, with diffusion coefficients of 0.08, 0.26, 0.84 μm$^2$/s and the same occupancy for each state (0.33). The state transition probabilities used for (A) were $p_{ii}$ = 0.8 and $p_{ij}$ = 0.1, which give rise to equal mass flow between each pairs of states; those used for (B) were $p_{12} = p_{23} = p_{31}$= 0.1 (counter-clockwise) and $p_{13} = p_{32} = p_{21}$ =0.8 (clockwise). Each simulation generated 5000 trajectories, which were then analyzed using vbSPT; each model was simulated 5 times, and the exemplary models with averaged model parameters are shown on the right. The resulting diffusion parameter outputs confirm that vbSPT was able to accurately determine parameters for both balanced (A, right) and non-balanced (B, right) state transitions. Error bars show 95% confidence interval.

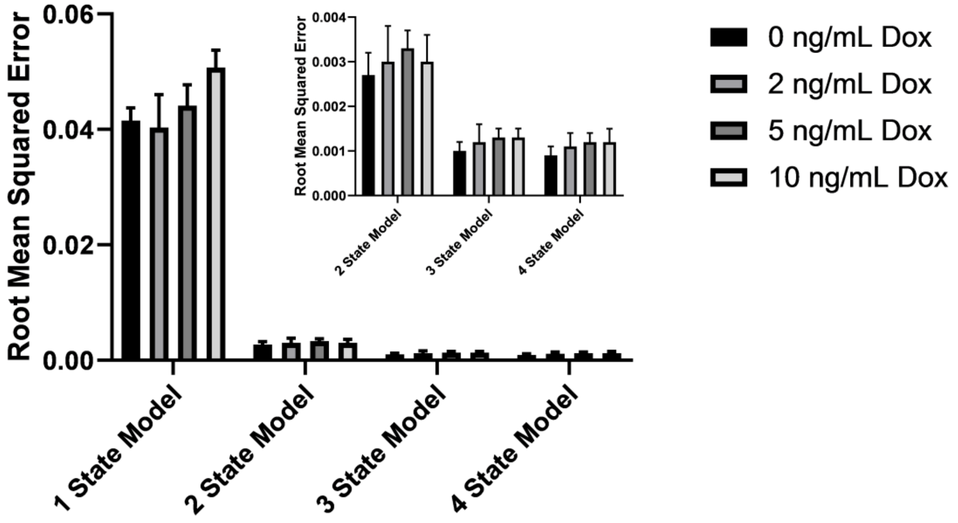


**Supplementary Figure 9. The three-state model remains optimal for KRas membrane diffusion over a broad range of expression levels.** The root mean squared error shown here is for CDF fitting of spt-PALM trajectories obtained at 0-10 ng/mL Dox, with all trajectories acquired at optimal conditions (<0.03 particles/μm$^2$ per frame and frame acquisition time 12 ms/frame). CDF fitting was used to fit data to one, two, three, and four state models, and the residual errors were calculated (as in Figure 1D).

#### References


1. Schmick, M. *et al.* KRas localizes to the plasma membrane by spatial cycles of solubilization, trapping and vesicular transport. *Cell* **157,** 459–471 (2014).

2. Lommerse, P. H. M. *et al.* Single-molecule diffusion reveals similar mobility for the Lck, H-Ras, and K-Ras membrane anchors. *Biophys. J.* **91,** 1090–1097 (2006).

3. Murakoshi, H. *et al.* Single-molecule imaging analysis of Ras activation in living cells. *Proc. Natl. Acad. Sci.* **101,** 7317–7322 (2004).
